# Genetic influences on hub connectivity of the human connectome

**DOI:** 10.1101/2020.06.21.163915

**Authors:** Aurina Arnatkevičiūtė, Ben D. Fulcher, Stuart Oldham, Jeggan Tiego, Casey Paquola, Zachary Gerring, Kevin Aquino, Ziarih Hawi, Beth Johnson, Gareth Ball, Marieke Klein, Gustavo Deco, Barbara Franke, Mark Bellgrove, Alex Fornito

**Author notes:** Contributed equally to this work (Ben D. Fulcher, Stuart Oldham).

## Abstract

Brain network hubs are both highly connected and highly inter-connected, forming a critical communication backbone for coherent neural dynamics. The mechanisms driving this organization are poorly understood. Using diffusion-weighted imaging in twins, we identify a major role for genes, showing that they preferentially influence connectivity strength between network hubs of the human connectome. Using transcriptomic atlas data, we show that connected hubs demonstrate tight coupling of transcriptional activity related to metabolic and cytoarchitectonic similarity. Finally, comparing over thirteen generative models of network growth, we show that purely stochastic processes cannot explain the precise wiring patterns of hubs, and that model performance can be improved by incorporating genetic constraints. Our findings indicate that genes play a strong and preferential role in shaping the functionally valuable, metabolically costly connections between connectome hubs.

## Introduction

Nervous systems are intricately connected networks with complex wiring patterns that are neither completely random nor completely ordered (1, 2). Numerous studies, conducted in species as diverse as the nematode *Caenorhabditis elegans*, mouse, macaque, and human, and at scales ranging from the cellular to the macroscopic, have shown that this complex organization is, in part, attributable to a heterogeneous distribution of connectivity across neural elements, such that a large fraction of network connections is concentrated on a small subset of network nodes called hubs (3–7). These hubs are more strongly interconnected with each other than expected by chance, forming a rich-club (3–5, 7) that is topologically positioned to integrate functionally diverse neural systems and mediate a large proportion of inter-regional communication (5, 8).

In human cortex, hubs are predominantly located in multimodal association areas (6, 9) and are among the most metabolically expensive elements of the connectome (10), with rich-club connections between hubs accounting for a disproportionate fraction of axonal wiring costs (3–5, 7, 11). Association hubs of the human brain also show marked inter-individual variability in connectivity and function that relates to a diverse array of behaviors (6, 12–14). These brain regions are disproportionately expanded in individuals with larger brains (15) and in human compared to non-human primates (16). They also show greater topological centrality and evolutionary divergence in the human connectome when compared to chimpanzee (17). These findings support the view that rapid expansion of multimodal association hubs, and the costly, valuable rich-club connections between them, underlies the enhanced cognitive capacity of humans compared to other species (18).

What influences the way in which hub regions connect to each other? The rapid evolutionary expansion of network hubs in humans, coupled with evidence supporting the heritability of many different aspects of brain organization (19), suggests an important role for genes. In the developing brain, neurons can innervate precise targets, even over long anatomical distances, by following genetically regulated molecular cues (20, 21). However, it is unknown whether genetic influences are preferentially exerted across specific classes of connections, such as the costly and functionally valuable links between network hubs. Preliminary evidence from human twin research suggests that certain properties of hub functional connectivity are strongly heritable (22), and analyses of *C. elegans*, mouse, and human data suggest that hub connectivity is associated with a distinct transcriptional signature related to metabolic function (7, 11, 23–25). Alternatively, some have suggested that the protracted maturation of hub regions (16, 26, 27) may endow these areas with enhanced plasticity (12), suggesting a prominent role for environmental influences. Moreover, recent computational models of whole-brain connectome wiring suggest that it is possible to grow networks with complex topological properties, including hubs, that mimic actual brains using simple, stochastic wiring rules based on geometric constraints (28–31) or trade-offs between the wiring cost and functional value of a connection (32– 34). These findings imply that the emergence of network hubs may not require precise genetic control and may instead result from random processes shaped by generic physical and/or functional properties.

Here, we use a multifaceted strategy to test between these competing views and characterize genetic influences on hub connectivity of the human cortical connectome. Using a connectome-wide heritability analysis (Fig. 1A-B), we show that genetic influences on phenotypic variance in connectivity strength are not distributed homogeneously throughout the brain, but are instead preferentially concentrated on links between network hubs. Then, as previously demonstrated in *C. elegans* (11) and mouse (7), we show that connected pairs of hubs in the human brain exhibit tightly coupled gene expression related to the metabolic demand and cytoarchitectonic similarity of these areas (Fig. 1C). Finally, we use computational modeling to show that stochastic network wiring models can indeed generate networks with brain-like properties, but fail to capture the spatial distribution of hub regions and, by extension, the precise pattern of wiring between network hubs. Moreover, adding genetic constraints to the models can improve their performance.

**Fig 1.**
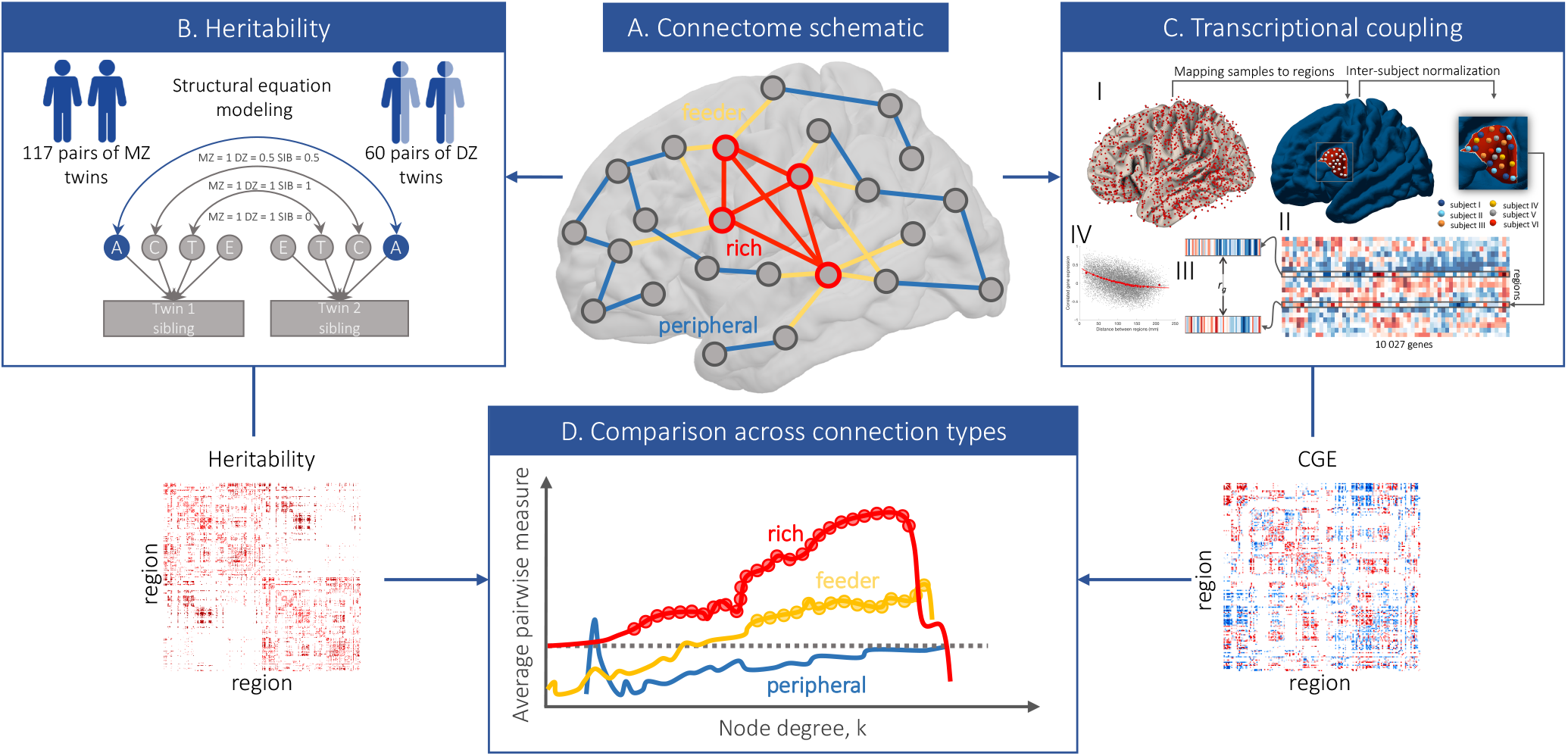
Workflows used to characterize genetic influences on hub connectivity. **(A)** A schematic representation of the connectome showing different connection types in the brain. Given a distinction between hub nodes (red outline) and nonhub nodes (grey outline), we can distinguish three classes of connections: rich links connections between two hubs (red); feeder links - connections between a hub and a nonhub (yellow); and peripheral links - connections between two nonhubs (blue). **(B)** Connectome-wide heritability analysis. We use structural equation modeling to fit a classic ACTE biometric model to every connection within the brain, resulting in estimates of genetic and environmental influences for each link. **(C)** Analysis of transcriptional coupling. (I) Each of 3702 tissue samples in the Allen Human Brain Atlas (AHBA) is mapped to a given region in our brain parcellation. (II) Expression values are then subjected to a quality control and processing pipeline (35) to construct a region × gene matrix of expression values. (III) We estimate correlated gene expression (CGE) between each pair of brain regions as the Pearson correlation between region-specific gene-expression profiles. (IV) Inter-regional CGE is corrected for spatial autocorrelation of the expression data via regression of an exponential distance trend (35). **(D)** Schematic representation of how values assigned to each edge are compared across connection types. We compare the mean of edge-level (pairwise) measures of heritability and CGE for rich, feeder, and peripheral links across all possible hub-defining thresholds (horizontal axis). As *k* increases, the definition of a hub becomes more stringent and identifies the actual hubs of the network. Thus, if a given effect is stronger for rich links, we expect the pairwise estimates to increase as a function of *k*, with the increase for rich links being particularly large relative to feeder and peripheral links.

Collectively, these findings demonstrate a direct link between molecular function and the large-scale network organization of the human connectome and highlight a prominent role for genes in shaping the costly and functionally valuable connections between network hubs.

## Results

Using diffusion weighted imaging (DWI) data for 972 subjects acquired through the Human Connectome Project (HCP) (36) we generate a representative group-level connectivity matrix [see Online methods] containing 12 924 unique connections between 360 brain regions defined by the HCPMMP1 atlas (37). This network contains a set of highly connected regions, quantified using the measure of node degree (*k*), which represent network hubs and span sensorimotor, paracentral, mid-cingulate (*k* > 105), insula, posterior cingulate, lateral parietal, and dorsolateral prefrontal cortices (*k* > 145) (Fig. 2A). As shown previously (5, 6, 8, 10), the network exhibits rich-club organization, with hubs being more densely and strongly interconnected than expected by chance (Fig. S1). Rich-club connections also have higher average wiring cost and communicability (Fig. S1), indicating that they are among the most topologically central and costly elements of the connectome.

**Fig 2.**
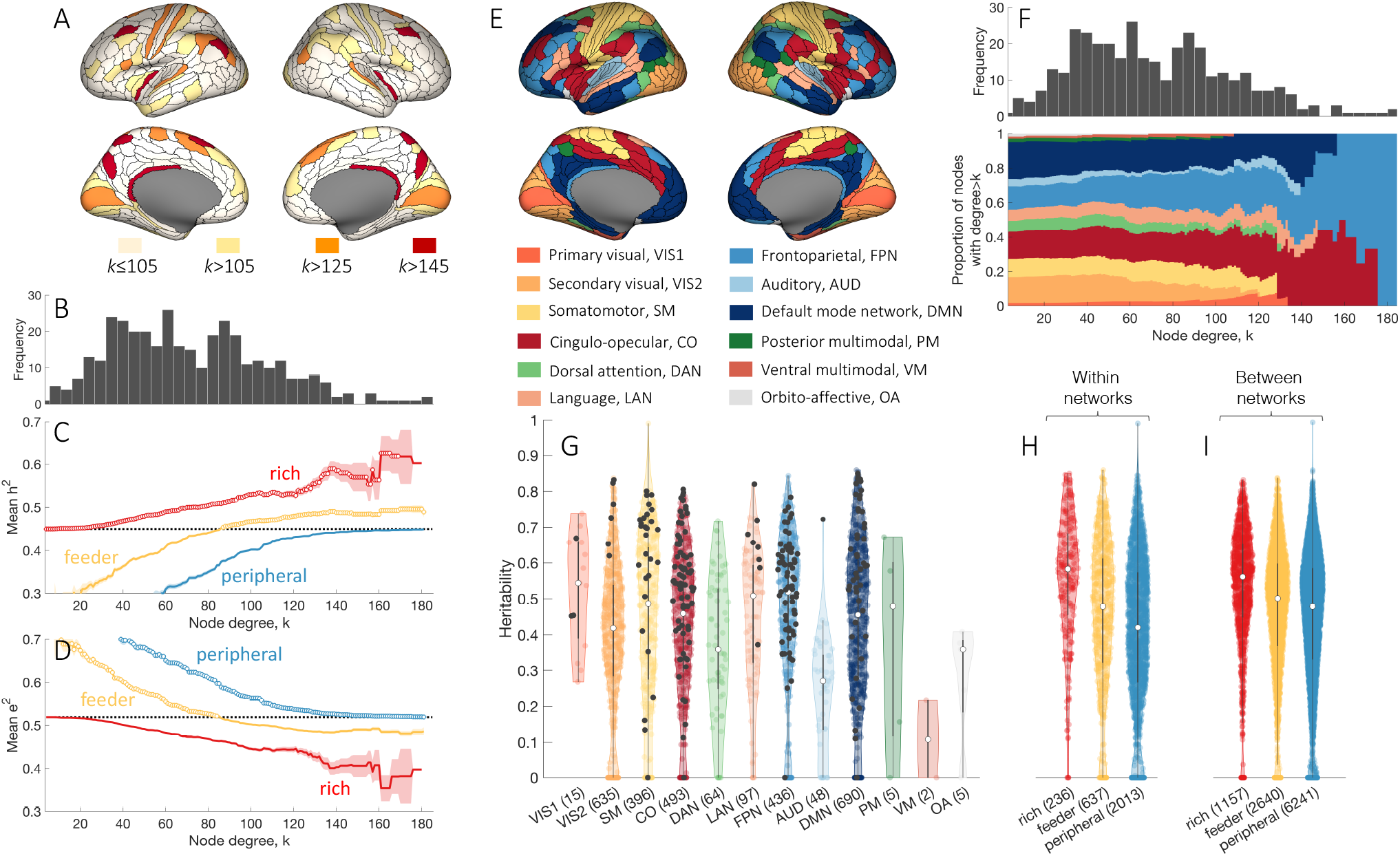
Genetic influences on connectivity strength are preferentially concentrated on rich-club links. **(A)** Anatomical locations of hubs defined at different levels of *k*. **(B)** The degree distribution of the representative group-level connectome. Mean genetic **(C)** and unique environmental **(D)** influences for rich (hub-hub), feeder (hub-nonhub), peripheral (nonhub-nonhub) connections as a function of the hub-defining threshold, *k*. The mean of the corresponding measure across all network links is shown as a dotted black line. Shaded area corresponds to the standard error of the mean, circles indicate a statistically significant increase of the measure in a given link type compared to the rest of the network (one-sided Welch’s *t*-test, uncorrected *p* < 0.05). **(E)** Regional assignments to canonical functional network modules (39), represented using color. **(F)** The proportion of nodes with degree > *k* in each functional network module as a function of *k*. **(G)** Distributions of heritability estimates across edges within functionally defined networks (39): VIS1 – primary visual; VIS2 – secondary visual; SM – somatomotor; CO – cingulo-opecular; DAN – dorsal attention; LAN – language; FPN – frontoparietal; AUD – auditory; DMN – default mode; PM – posterior multimodal; VM – ventral multimodal; OA – orbito-affective. Rich links within each module are represented as black dots, as defined for *k* > 105. Heritability distributions for edges within **(H)** and between **(I)** functional modules across rich, feeder, and peripheral link types for *k* > 105. Rich links show significantly higher heritability compared to both feeder and peripheral links, within and between functional modules (one-sided Welch’s *t*-test, all *p* < 1.9 × 10^−12^).

### Genetic influences on brain connectivity are concentrated in the rich club

To investigate whether genes preferentially influence certain classes of connections in the human brain, we perform a connectome-wide heritability analysis of twin data acquired through the Human Connectome Project. For each of 234 monozygotic (MZ) twins and their 69 non-twin siblings as well as 120 dizygotic (DZ) twins and 48 of their non-twin siblings, we reconstruct macroscale cortical connectomes using DWI [see Online methods].

For each connection in the representative group connectome, we use the classic ACTE model to estimate the proportion of variance in connectivity strength that is attributable to additive genetic factors (narrow-sense heritability, denoted *h*^2^, see Online methods). Using the average fractional anisotropy (FA) of each connecting fiber bundle to quantify connectivity strength [see Online methods], we observe a wide range of heritability estimates across connections, spanning 0 to 0.99 (*h*^2^_mean_ = 0.45, *h*^2^_SD_ = 0.2). Non-trivial genetic influences, quantified using the A component of the ACTE model, are observed for the majority of connections, with the AE model showing the best fit for 86.7% of edges, ACTE for 4.3%, and ACE for 1.3%. A total of 7.7% of connections are influenced only by environmental factors (CE model 6.8%, E model 0.9%). To examine whether genetic influences are preferentially concentrated on specific types of inter-regional connections, we distinguish between hub and nonhub regions, resulting in three possible types of connections: rich (hub-to-hub), feeder (between a hub and a nonhub), and peripheral (nonhub-tononhub) links [see schematic in Fig. 1A, (38)]. We find that mean heritability derived from the best-fitting biometric model is highest for rich, intermediate for feeder, and lowest for peripheral connections across nearly all values of *k* (Fig. 2B–C). The increase in heritability for rich links as a function of the hub-defining threshold, *k*, indicates that genetic influences are, on average, stronger for connections between the most highly connected brain regions (see also Fig. 1D). The same pattern is replicated when taking genetic parameters from the full ACTE model, confirming that this result is not an artefact of our model-selection procedure (Fig. S2B).

Contrasting with rich links between hubs, phenotypic variance in peripheral connectivity (between nonhubs) is predominantly influenced by unique environment (quantified by the model parameter E, Fig. 2D), while common environmental influences are consistently low across all link types (mean values ⟨*C*⟩ < 0.08 and ⟨*T*⟩ < 0.02 across all *k* thresholds). Critically, we obtain similar evidence of preferential genetic influences on hub connectivity when using different methods for parcellating or thresholding our connectomes (Fig. S3) or when evaluating connection strength based on the number of reconstructed streamlines (streamline count, SC) between regions (Fig. S2C,D).

To investigate whether genetic influences are specific to certain functional systems of the brain, we next categorize edges according to the major functional networks that they connect, as defined using a network parcellation (39) of the HCPMMP1 atlas (37) (Fig. 2E). Figure 2F shows the proportion of nodes with degree > *k* in each functional network. High-degree nodes are present in most networks until *k ≈* 120, beyond which they are predominantly found in multimodal association networks; namely, the fronto-parietal, cinguolo-opercular, and default mode systems.

Across all 12 canonical functional networks, rich links both within and between networks demonstrate significantly higher heritability than other types of connections (Fig. 2G-I, one-sided Welch’s *t*-test, comparing heritability of rich *vs* feeder and rich *vs* peripheral connections, all *p* < 1.9 × 10^−12^), indicating that the elevated genetic influences observed for rich links cannot be explained by the affiliation of hub nodes to any specific functional network. Moreover, stronger heritability of rich links is evident across different connection distances (Fig. S4A-C), suggesting that preferential genetic influences on hub connectivity cannot be explained simply by the longer average distance of rich links (Fig. S1D). Other factors, such as the number of outliers excluded from the analysis (Fig. S4D-F) or differences in the phenotypic variance of connectivity strength estimates across different edge types (Fig. S4G-I), were also unable to account for the increased heritability of rich links.

Together, these findings indicate that genetic influences on phenotypic variance in connectivity strength are not distributed homogeneously throughout the brain, nor are they confined to specific functional networks or long *vs* short-range connections. Instead, they are most strongly concentrated on the connections between network hubs. These hubs are distributed throughout the cortex, with the most highly connected regions residing in multimodal association networks.

### Transcriptional coupling is elevated between connected hubs

Next, we investigate the transcriptional correlates of hub connectivity using data from the Allen Human Brain Atlas (AHBA) (40), focusing on expression profiles of 10 027 genes surpassing our quality-control criteria (35) within the 180 cortical regions of the left hemisphere, where spatial coverage in the AHBA is most comprehensive. Evaluating CGE across the full set of genes allows us to quantify the global expression patterns across the brain without restricting the analysis to predefined gene categories. We secondarily test for enrichment of certain classes of genes, as detailed below. We quantify transcriptional coupling between different brain regions using spatially-corrected correlated gene expression (CGE) (Fig. 1C, Fig. S5, see also Online methods) and define inter-regional connectivity using a binary group-representative matrix [see Online methods]. The spatial correction is important as prior studies of *C. elegans*, mouse, and human nervous systems have shown that, across the brain, CGE decays exponentially as a function of distance (7, 11, 24, 41). Recent analyses of the mesoscale connectome of the mouse (7) and microscale (cellular) connectome of *C. elegans* (11) indicate that, after considering this bulk trend, connected pairs of hubs show the highest CGE, despite being separated by longer anatomical distances, on average, than other neural elements.

Figures 3A–B show that the same effect is observed in humans: CGE is highest for rich, intermediate for feeder, and lowest for peripheral links. We obtain similar results under different connectome processing options (Fig. S6), distance ranges (Fig. S7), and when using connectivity data from an independent sample (Fig. S8A). The consistency of this effect between human, mouse, and *C. elegans* [see Fig. S9 for comparison] is striking given the large physiological differences between species, methods for connectome reconstruction (DWI, viral tract tracing, electron microscopy), analysis resolution [macroscale (mm to cm), mesoscale (*µ*m to mm), microscale (individual cells and synapses)], and gene-expression assays (microarray, *in situ* hybridization, curation of published reports).

**Fig 3.**
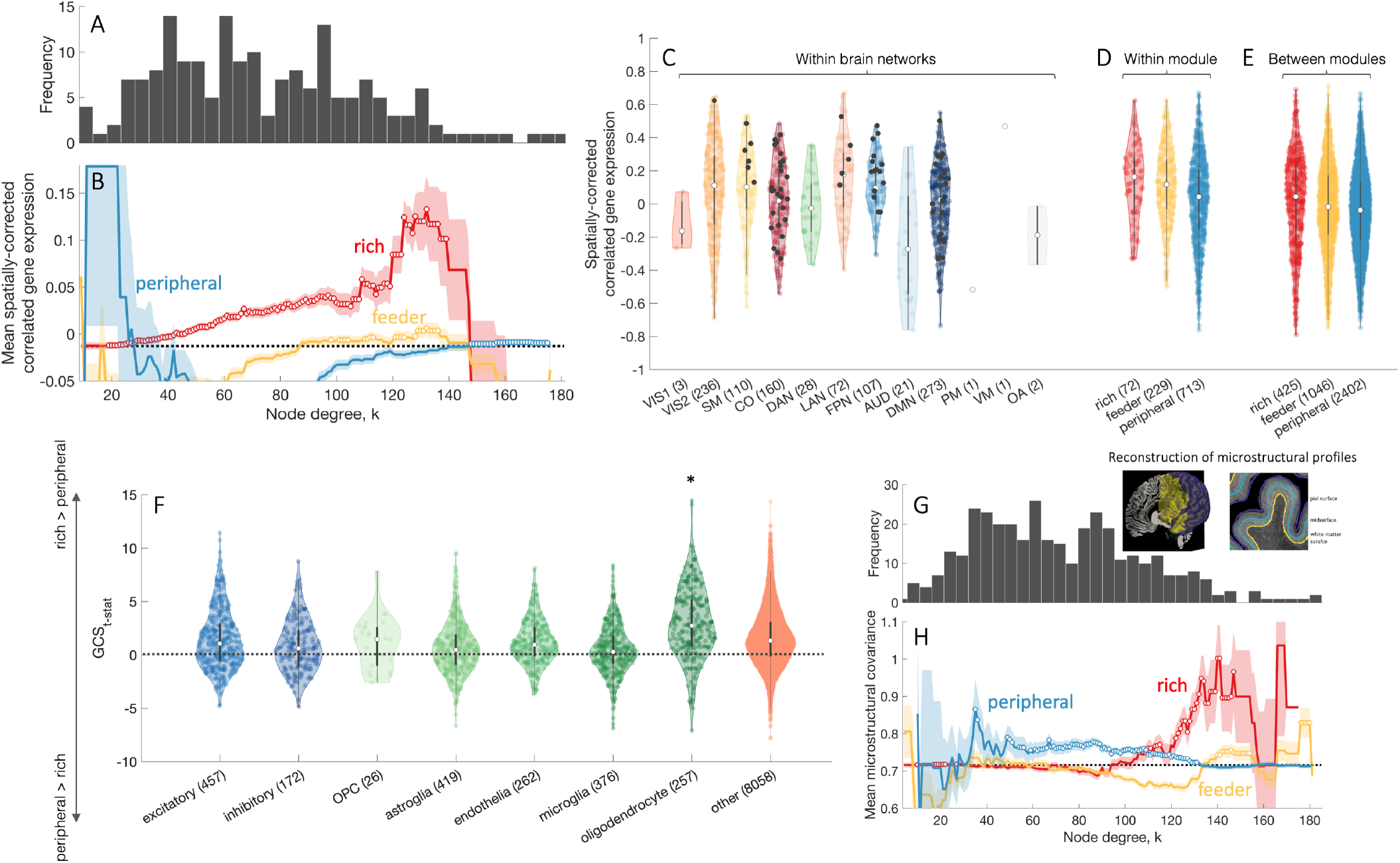
Transcriptional coupling is elevated for connected brain network hubs. **(A)** The degree distribution of the representative group-level connectome of brain regions in the left cortical hemisphere. Degree is computed from whole-brain connectivity. **(B)** Mean correlated gene expression (CGE) for rich (hub-hub), feeder (hub-nonhub), peripheral (nonhub-nonhub) connections as a function of the degree threshold, *k*, used to define hubs. The mean CGE across all network links is shown as a dotted black line. The shaded area corresponds to the standard error of the mean, circles indicate a statistically significant increase in CGE in a given link type compared to the rest of the network (one-sided Welch’s *t*-test, uncorrected *p* < 0.05). CGE estimates are corrected for distance effects, as explained in the Online methods. **(C)** CGE within functionally defined networks as in Fig. 2E. Black dots represent CGE values for rich links (*k* > 105). CGE values within **(D)** and between **(E)** functional modules in the left hemisphere across different link types (rich, feeder, and peripheral). Inter-module rich links show significantly higher CGE compared to both feeder (one-sided Welch’s *t*-test, *p* = 0.03) and peripheral links (*p* = 1.5 × 10^−4^). Within functional modules, rich links show higher CGE compared to peripheral (*p* = 1.2 × 10^−4^) but not to feeder links (*p* = 0.5). **(F)** Gene contribution score *t*-statistic values (GCS_t−stat_) for cell-specific gene groups quantifying the contribution of individual genes towards increased CGE for rich compared to peripheral links. Neuronal gene groups (excitatory – excitatory neurons; inhibitory – inhibitory neurons) are colored blue; glial gene groups (OPC – oligodendrocyte progenitor cells, astroglia, endothelia – endothelial cells, microglia, oligodendrocytes) colored green; values for all other genes presented in light orange. Oligodendrocyte-related genes show a statistically significant increase in GCC compared to all other genes (one-sided Welch’s *t*-test, *p* = 2 × 10^−11^). **(G)** The degree distribution of the representative group-level cortical connectome. **(H)** Mean microstructural profile covariance (MPC) for rich (hub–hub), feeder (hub–nonhub), peripheral (nonhub–nonhub) connections as a function of degree threshold, *k* used to define hubs. The MPC across all network links is shown as a dotted black line. Shaded area corresponds to the standard error of the mean, circles indicate a statistically significant increase in MPC in a given link type compared to the rest of the network (one-sided Welch’s *t*-test, uncorrected *p* < 0.05). Inset near the degree distribution shows examples of the intermediate surfaces used to assay microstructure across the cortical depth.

As with heritability (Fig. 2C), higher CGE occurs for connections between high-degree nodes distributed across the brain; i.e., the effect is not confined to a single functional network (Fig. 3C). Indeed, connected pairs of hubs demonstrate higher CGE both within (Fig. 3D) and between (Fig. 3E) functionally defined networks (onesided Welch’s *t*-test, comparing CGE of rich *vs* feeder and rich *vs* peripheral connections, all *p* ≤ 0.02).

Expression values in the AHBA are extracted from bulk tissue samples, and thus agglomerate transcriptional information from many different cell types. It is therefore possible that inter-regional CGE may be related to similarity in regional cellular composition [see also (42)]. We thus repeated the CGE analysis, this time using only data from genes showing cell-specific expression for seven canonical cell types: excitatory and inhibitory neurons, oligodendrocyte progenitor cell, astroglia, endothelial cells, microglia, and oligodendrocytes [see Online methods, (43–47)]. We find that all classes of cellspecific genes exhibit an increase in CGE for rich links relative to peripheral (Fig. S10), with oligodendrocyte-related genes showing a significantly stronger contribution to elevated CGE between connected hubs (one-sided Welch’s *t*-test, *p* = 2 × 10^−11^, Fig. 3F) compared to all other genes [see Online methods].

These findings suggest that connected hubs may have higher cytoarchitectonic similarity than other pairs of regions. Given that the CGE of cell-specific genes is a relatively indirect marker of cytoarchitecture, we conducted a more direct test of the hypothesis that connected hubs have more similar cytoarchitecture using the BigBrain atlas (48), which is a high-resolution Merker-stained histological reconstruction of a post-mortem human brain that provides an opportunity to map regional variations in cellular density as a function of cortical depth. Following Paquola et al. (49), we estimate intensity profiles across 16 equivolumetric surfaces placed between the gray/white and pial boundaries of the cortical ribbon and compute the inter-regional microstructural profile covariance (MPC) [see Online methods] as a proxy for cytoarchitectonic similarity. Mirroring the CGE and heritability findings, rich links exhibit elevated MPC compared to feeder and peripheral edges (Fig. 3H). These convergent MPC and cell-specific CGE results indicate that connected hubs have a more similar cytoarchitecture than other pairs of brain regions.

Finally, a gene set enrichment analysis of gene groups related to elevated CGE between hubs (Online methods) identifies significant enrichment of 49 GO categories, notably featuring genes related to oxidative metabolism, ATP synthesis coupled electron transport, and mitochondrial function (*p*_FDR_ < 0.05, Table S1) that mirror those previously reported in the mouse brain (7). These results suggest a close genetic link between hub connectivity and metabolic function [for additional considerations see (50) and Online methods].

### Stochastic models of brain wiring do not capture the spatial distribution of degree

The results above imply strong genetic control of hub connectivity, which seems at odds with recent modeling studies suggesting that simple stochastic wiring rules can generate networks with complex, brain-like topologies, including heavy-tailed degree distributions that signify the existence of hubs (29, 30, 32, 33). Investigating variability in the binary topology of connectivity–that is, the specific pattern of wiring between regions–is challenging, as such variations at the level of macroscale human connectomes are limited. It is thus possible that stochastic processes may give rise to the basic binary topology of hub connectivity, with variations in connectivity strength subsequently being influenced by genetic factors.

To investigate the role of stochastic processes in shaping hub connectivity, we fitted 13 different generative models of network wiring to the HCP connectome data. Under each model, synthetic connectomes are generated using probabilistic wiring rules. The models we consider here have been explored extensively in prior work (33) and have the general form:

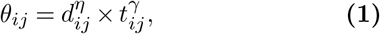

where *θ*_*ij*_ is a score that weights the probability of connecting nodes *i* and *j, d*_*ij*_ is the Euclidean distance between node pairs, and *t*_*ij*_ is a topological property of an edge that may confer functional value to the network. Each of the 13 models substitutes a different topological property for *t*_*ij*_ (definitions in Table 2). The exponents *η* and *γ* are free parameters fitted to the data to optimally match the topological properties of the actual human connectome, as defined using nodal distributions of degree, clustering, and betweenness, and the edge-level distribution of connection distances [(33), see Online methods].

In line with prior work (33), we find that models in which connections form according to both spatial (wiring cost) and topological rules can fit the distributions of empirical network properties better than a model based on wiring cost alone (i.e., the ‘sptl’ model), as shown in Fig. 4A. The best-fitting model, ‘deg-avg’, modulates a pure wiring cost term by favoring connectivity between pairs of nodes that already have high average degree, and shows a good fit to the data (i.e., all fits, indexed by the Kolmogorov-Smirnov statistic, were KS < 0.21 [see Online methods for an extended discussion]).

**Fig 4.**
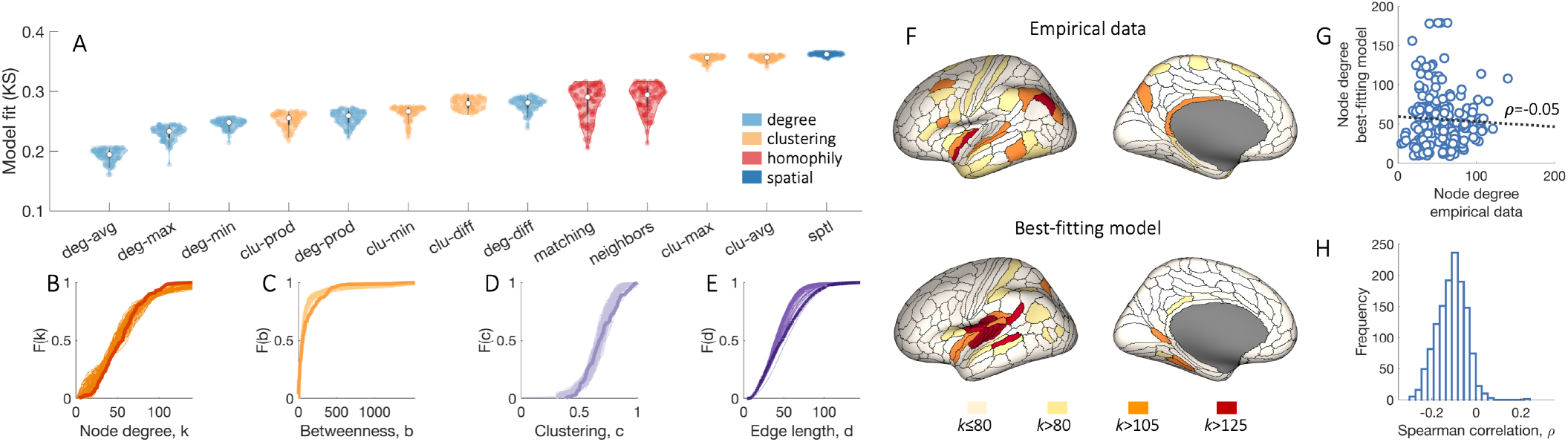
Generative brain network models do not reproduce the spatial distribution of brain network hubs. **(A)** Each distribution represents estimates of model fit, as quantified by the maximum KS value of the top 100 networks (out of 10 000) produced by the model optimization procedure. The color of each box indicates conceptually related models, as determined by the specific topology metric used in the model [see Table 2]. Models favoring homophilic connectivity between node pairs are shown in red, those favoring clustering in orange, those based on degree in light blue, and a purely spatial model considering wiring costs alone is in dark blue. The specific wiring-rule names are shown along the horizontal-axis, with formal definitions provided in Table 2. Cumulative distributions of: **(B)** node degree, *k*; **(C)** betweenness centrality, *b*; **(D)** clustering coefficient, *c*; and **(E)** edge length, *d*, for the empirical connectome (darker line) and 100 runs (lighter lines) of the best-fitting ‘deg-avg’ model corresponding to the data points shown in **A. (F)** Anatomical locations of hubs defined for a single hemisphere at selected *k* thresholds for the empirical data (top) and the single run of the optimized ‘deg-avg’ generative model demonstrating the best model fit across 10 000 runs (bottom). **(G)** Correlation between the degree sequences of the empirical data and the best-fitting generative model within a single hemisphere (Spearman’s *ρ* = −0.05). **(H)** The distribution of correlation values quantifying the relationship between left hemisphere degree sequences of the empirical data and synthetic networks generated using the top 100 best-fitting parameter combinations for each of the 13 considered models, corresponding to the data points shown in **A**.

Despite this adequate fit to four key network properties of the human connectome (Figs. 4B–E), we find that node degree in the empirical and model networks have very different spatial distributions. As shown in Fig. 4F, hubs in the empirical data are distributed throughout the brain, whereas hubs in the network that demonstrates the best fit to data across 130 000 model runs are predominantly confined to temporal cortex. As a result, the correlation between the degree sequences of the empirical and model networks is very low (e.g., *ρ* = − 0.05, Fig. 4G). This low correlation is observed consistently across all models (Fig. 4H), and even when we fit model parameters to explicitly optimize the correlation between empirical and model degree degree sequences [see Fig. S11, Online methods]; across 260 000 model runs, the degree sequence correlation with the empirical data never exceeds *ρ* = 0.3.

Together, these findings indicate that while stochastic models of brain network wiring can capture the statistical properties (node- and edge-level distributions) of connectomes, they cannot reproduce the way in which these properties are spatially embedded and thus do not accurately replicate the precise pattern of wiring between connectome hubs.

### Genetically constrained models offer improved fits to topological and topographical properties of the connectome

The limitations of stochastic models, coupled with our evidence for a strong genetic influence on hub connectivity, raises the question of whether models that include genetic information may show better performance than models based on cost and/or topology alone. To address this question, we focus on the best-fitting cost-topology model, ‘deg-avg’, and examine its performance relative to model variants that include a bias to form connections between pairs of regions with high CGE affects. We focus specifically on CGE, given our evidence for elevated CGE between pairs of hubs regions (Fig.3).

Figure 5 compares model fit statistics for the original ‘deg-avg’ model (denoted ‘ST’ in Fig.5A), and models in which connections are formed according to CGE alone (denoted ‘G’), wiring cost alone (denoted ‘S’), an interplay between CGE and wiring cost (denoted ‘SG’), and an interplay between CGE and topological constraints, as defined in the ‘deg-avg’ model (denoted ‘TG’). We find that a model incorporating both topology and genetic information, such that connections are favored between regions with both high average degree and high CGE (‘TG’ model), shows the best fits, on average, to network topology, even surpassing the ‘deg-avg’ (‘ST’) model.

**Fig 5.**
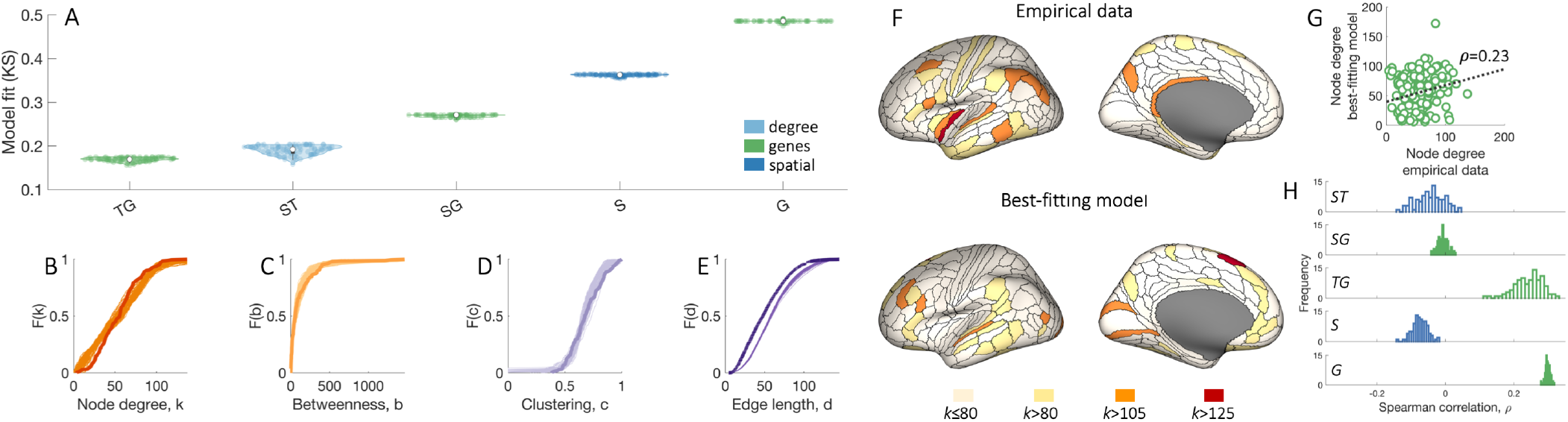
Adding genetic constraints to generative models can improve fits to network topology and topography. **(A)** Each distribution represents estimates of model fit, as quantified by the maximum KS value of the top 100 networks (out of 10 000) produced by the model optimization procedure. The color of each box indicates conceptually related models, as determined by the specific metric used in the model: models favoring connectivity between regions with similar gene expression are in green, a model based on degree and wiring cost is in light blue, and a purely spatial model considering wiring costs alone is in dark blue. ‘S’, ‘T’, ‘G’ stand for space (wiring cost), topology and gene expression respectively. Cumulative distributions of: **(B)** node degree, *k*; **(C)** betweenness centrality, *b*; **(D)** clustering coefficient, *c*; and **(E)** edge length, *d*, for the empirical connectome (darker line) and 100 runs (lighter lines) of the best-fitting ‘TG’ model corresponding to the data points shown in **A. (F)** Anatomical locations of hubs defined for a single hemisphere at selected *k* thresholds for the empirical data (top) and the single run of the optimized ‘TG’ generative model demonstrating the best model fit across 10 000 runs (bottom). These networks contain 177 regions (instead of 180 presented in Fig.4F) due to the limited coverage of gene expression data. **(G)** Correlation between the degree sequences of the empirical data and the best-fitting generative model within a single hemisphere (Spearman’s *ρ* = 0.23). **(H)** The distributions of correlation values quantifying the relationship between left hemisphere degree sequences of the empirical data and synthetic networks generated using the top 100 best-fitting parameter combinations for each of the 6 considered models, corresponding to the data points shown in **A**.

Moreover, the spatial distribution of hubs in the ‘TG’ model network demonstrating the best fit is more dispersed across the brain (Fig. 5F) compared to the classical ‘deg-avg’ model, resulting in a higher correlation between degree sequences of the empirical and model networks (Fig. 5G). Overall, models including topology and CGE or CGE alone demonstrate more positive degree sequence correlations compared to models that do not include CGE (Fig. 5H). Our findings thus indicate that incorporation of genetic constraints into stochastic models of the connectome can improve fits to both network topology and spatial topography.

## Discussion

The complex topology of neural networks is thought to have been sculpted by competitive selection pressures to minimize wiring costs and promote complex, adaptive function (10, 51). Across diverse species, rich-club connections between hubs are among the most costly and topologically central links of the connectome (3–5, 7) and thus play a major role in determining how cost–value trade-offs are negotiated within a given nervous system. Here, we combine a multifaceted genetic analysis with mathematical modeling to examine the mechanisms that shape hub connectivity of the human connectome. We find that: (i) genetic influences on phenotypic variation in connection strength are principally concentrated on the rich links between hubs; (ii) connected hubs have highly correlated gene expression patterns that are related to similarity in regional cytoarchitecture and energy metabolism; (iii) current stochastic models of network growth cannot reproduce the spatial distribution of hubs; and (iv) adding genetic constraints to these models can improve performance. Together, these findings support a major role for genes in shaping the rich-club organization of the brain.

Our connectome-wide heritability analysis presents evidence for a non-uniform distribution of genetic influences across the brain, characterized by a gradient in which genetic influences are weak for peripheral connections between nonhubs, intermediate for feeder connections between hubs and nonhubs, and strongest for rich links between hubs. Critically, this effect cannot be attributed to connection distance or network affiliation, suggesting some degree of specificity to hubs located throughout the brain.

The most strongly connected hubs in our connectomes were located in multimodal association networks, which show disproportionate expansion in size and connectivity in human compared to nonhuman primates (16–18). Given the high centrality and cost of these connections [Fig. S1, (3–5, 7)], the preferential genetic influence on rich-club connectivity that we observe supports the hypothesis that natural selection favors wiring patterns that provide high value for low cost and that selection pressures are strongly concentrated on the valuable, costly links between hubs (2, 10). This view is also supported by recent evidence that genes demonstrating accelerated divergence between humans and chimpanzees show elevated expression in multimodal association networks (52).

In contrast to rich-club connectivity, phenotypic variance in peripheral connections between nonhubs is primarily influenced by unique environment. Topologically peripheral connections are more strongly conserved between human and chimpanzee connectomes (17). Moreover, the spatial topography and function of nonhub sensory areas is highly consistent across primates, presumably being specified early in development by evolutionarily conserved transcriptional gradients (18). These conserved gradients may couple with simple physical processes to give rise to predominantly short-range connectivity between topologically peripheral pairs of regions (31, 53). Subsequent modifications to peripheral connectivity may be driven by activity-dependent mechanisms, resulting in a greater environmental influence on phenotypic variance in connection strength.

It has been proposed that evolutionary expansion of multimodal hubs untethers these regions from transcriptional anchors in sensory areas, resulting in distinctive, non-canonical anatomical and functional properties (18). Our findings suggest that, despite this putative untethering, genes still play an important role in shaping phenotypic variance of hub connectivity. This result aligns with evidence that non-conserved network properties reflect evolutionary innovations that are driven by structural variation of DNA, yielding greater phenotypic variation within a species (12) and higher trait heritability when compared to more strongly conserved properties (54).

In addition to being highly heritable, pairs of connected hubs also show the highest levels of transcriptional coupling, as previously observed in the mesoscale mouse connectome (7) and cellular connectome of *C. elegans* (11) [see Fig. S9]. In *C. elegans* this result is not explained by the developmental proximity (i.e., similarity in neuron birth time or cell lineage distance), neurochemical identity, or anatomical position of neuron pairs, but is instead related to the functional identity of hub neurons, which tend to be command interneurons. Our analysis of cell-specific genes suggests a similar result at the regional level in humans, as rich-link CGE was elevated for gene markers of seven different cell types suggesting that network hubs have enhanced similarity in regional cytoarchitecture. This conclusion was supported by our MPC analysis of the BigBrain atlas. Our results align with the structural model of cortical connectivity, in which regions with similar cytoarchitecture are more likely to connect with each other, even over long distances (55). More specifically, our findings suggest that hub areas are the most similar in their cellular composition, and that this similarity may play a critical role in how genes preferentially sculpt long-range interconnectivity between hubs.

We also show that current stochastic models of network growth, despite capturing key statistical network properties of the connectome, do not reproduce the spatial locations of network hubs. Indeed, while the actual hubs of the human brain have a widespread anatomical distribution, hubs in the best-fitting (deg-avg) model network are concentrated around centrally located regions. In line with this result, recent work has shown that cost-neutral randomizations, in which connections are progressively randomized while preserving total wiring cost and the existence (but not position) of hubs, almost always degrade the functional complexity of the network, disconnect high-cost hubs, and lead to a distinct hub topography in which the most highly connected nodes cluster near the centre of the brain (56). These findings suggest that actual brains are very close to optimally balancing wiring cost with topological complexity, and that hub connectivity plays a critical role in determining how this balance is realized [see also (57)].

Notably, we find that incorporating genetic constraints into the models improves their capacity to reproduce both network topology and the spatial topography of hubs. While there is still room for considerable further improvement, our findings indicate that combining topological, genetic, and possibly spatial information may offer a fruitful way forward for generative models of the human connectome. Indeed, some models suggest that random growth of connections, when coupled with changes in brain geometry and heterochronicity of connection formation across regions, can yield brainlike networks with realistic features (29), including connectivity between regions with similar cytoarchitecture (58, 59). Although genes likely influence heterochronous development, future work extending such models so that they can be directly fitted to empirical data in humans, and considering which genes may be most relevant in shaping network wiring, will be important for delineating the precise roles of genetic, environmental, stochastic, and physical mechanisms in shaping connectome architecture.

## Online methods

Code for reproducing the results presented here is available at github https://github.com/BMHLab/GeneticBrainHubs. Data are available from an associated figshare repository.

### Imaging data acquisition

We examined DWI data from two independent cohorts. The first was obtained from the Human Connectome Project [HCP, (36)]. We used the minimally processed DWI and structural data from the HCP for 972 participants (age mean ± standard deviation: 28.7 ± 3.7, 522 females), including a cohort of MZ and DZ twin pairs together with their non-twin siblings (more details presented in Online methods). Data were acquired on a customized Siemens 3T “Connectome Skyra” scanner at Washington University in St Louis, Missouri, USA using a multi-shell protocol for the DWI: 1.25 mm^3^ isotropic voxels, repetition time (TR) = 5520 ms, echo time (TE) = 89.5 ms, field-of-view (FOV) of 210 × 180 mm, 270 directions with b = 1000, 2000, 3000 s/mm^2^ (90 per b value), and 18 b = 0 volumes. Structural T1-weighted data were collected using 0.7 mm^3^ isotropic voxels, TR = 2400 ms, TE = 2.14 ms, FOV of 224 × 224 mm. The full details can be found elsewhere (60).

The second DWI dataset came from individuals recruited as part of ongoing research conducted at Monash University. This Monash sample was used for replication of the CGE analysis, and comprised 439 participants with MRI data obtained on a Siemens Skyra 3T scanner at Monash Biomedical Imaging in Clayton, Victoria, Australia using the following parameters: 2.5 mm^3^ voxel size, TR = 8800 ms, TE = 110 ms, FOV of 240 × 240 mm, 60 directions with b = 3000 s/mm^2^ and seven b = 0 volumes. In addition, a single b = 0 s/mm^2^ was obtained with the reverse-phase encoding so distortion correction could be performed. T1-weighted structural scans were acquired using: 1 mm^3^ isotropic voxels, TR = 2300 ms, TE = 2.07 ms, FOV of 256 × 256 mm. Data for 15 subjects were excluded due to: low connectome density (*n* = 10, connectome density more than 3 standard deviations lower than the mean) or issues with cortical surface segmentation (*n* = 5), resulting in a final sample of 424 participants (age mean ± standard deviation: 23.5 ± 5.3, 190 females).

### Image pre-processing

HCP DWI data were processed according to the HCP minimal preprocessing pipeline, which included normalization of mean *b*_0_ image across diffusion acquisitions, and correction for EPI susceptibility and signal outliers, eddy-current-induced distortions, slice dropouts, gradient-nonlinearities and subject motion. T1-weighted data were corrected for gradient and readout distortions prior to being processed with Freesurfer [full details can be found in (60)].

Pre-processing for T1-weighted structural images in the Monash Sample consisted of visual screening for gross artefacts followed by the reconstruction of the grey/white matter interface and the pial surface using FreeSurfer v5.3.0 software. Surface reconstructions for each subject were visually inspected, with manual corrections performed as required to generate accurate surface models (60).

Distortions in the Monash DWI data were corrected with TOPUP in FSL, using the forward and reverse phase-encoded *b* = 0 images to estimate the susceptibility-induced off-resonance field (61, 62). We corrected for eddy-current distortions, volume-to-volume head motion, within-volume head motion, and signal outliers using eddy tool in FSL [version 5.0.11; (63–65)]. This implementation of EDDY significantly mitigates motion-related contamination of DWI connectivity estimates (66). DWI data were subsequently corrected for B1 field inhomogeneities using FAST in FSL (62, 67).

### Connectome reconstruction

For both the HCP and Monash datasets, network nodes for each individual were defined using a recently-developed, data-driven group average HCPMMP1 parcellation of the cortex into 360 regions [180 per hemisphere, (37)]. An advantage of this parcellation is that it uses diverse structural and functional information to derive a consensus partition of the cortex into different areas. Each region has also been assigned to a distinct canonical functional network (39), allowing us to examine results in relation to the organization of these classic systems. However, the resulting areas can vary considerably in size, which can affect regional connectivity estimates since larger regions are able to accommodate more connections. To ensure that our results were not driven by the use of this specific parcellation, we replicated our main findings using a random cortical parcellation consisting of 500 approximately equally sized regions [250 per hemisphere, generated using the approach described in (68); code available at https://github.com/miykael/parcellation_fragmenter]. This offers a stringent test of the generalizability of our findings, as the parcellations vary in terms of both method for construction (data-driven *vs* random) and resolution (360 *vs* 500 nodes).

We focus our analysis on cortical connectivity for simplicity, as we know of no unified parcellation of cortical and subcortical areas, positional differences between cortex and subcortex can affect DWI connectivity estimates, and there are major differences between cortical and subcortical patterns of gene expression in the AHBA data that can be difficult to appropriately accommodate (35).

Subsequent processing of the DWI data for both the HCP and Monash data was performed using the MRtrix3 (69) and FMRIB Software Library (70). Tractography was conducted in each participant’s T1 space using second order integration over fibre orientation distributions (iFOD2) (71). To further improve the biological accuracy of the structural networks, we also applied Anatomically Constrained Tractography (ACT), which uses a tissue segmentation of the brain into cortical grey matter, subcortical grey matter, white matter, and cerebrospinal fluid to ensure that streamlines are beginning, traversing, and terminating in anatomically plausible locations (72). Tissue types were determined using FSL software (70). A total of 10 million streamlines were generated on a probabilistic basis using a dynamic seeding approach that evaluates the relative difference between the estimated and current reconstruction fibre density and preferentially samples from areas of insufficient density (73). This method helps mitigate biases related to poor reconstruction of tracts from certain parts of the brain due to insufficient seeding. The resulting tractogram was then combined with the cortical parcellation for each subject to produce a network map of white matter connectivity. Streamline termination points were assigned to the closest region within a 5 mm radius.

Connection weights were quantified using both streamline count (number of streamlines connecting two regions, SC) and the mean fractional anisotropy (FA) of voxels traversed by streamlines connecting two regions, which is commonly used as a marker of white matter microstructure. We focused on SC and FA as measures of connectivity strength because they are the most widely used in the literature, but note that they can be influenced by numerous factors that are not directly related to physiological measures of communication capacity between two regions (74). Moreover, while diffusion tractography remains the only available tool for *in vivo* connectivity mapping in humans, tractography algorithms can vary in their specificity and sensitivity for tract reconstruction (75, 76). To mitigate these effects, our data processing pipeline has been designed to limit contributions from spurious streamlines (72) and head motion (66). While the accuracy of all tractography methods remains an open challenge for the field (77), we note that any errors in tract reconstruction should reduce our chances of identifying stronger genetic effects for rich links through a heritbility analysis, since noisy connectivity values will inflate estimates of the E parameter (which also accounts for measurement error) in our biometric models, and rich links tend to be long-range connections, which are more prone to tractography errors (78). Our findings may thus provide a conservative estimate of genetic influences on hub connectivity.

### Connectome thresholding

As connectomes are estimated with some degree of noise, it is common practice to threshold weak or inconsistent edges to focus on connections that can be more reliably estimated (79). We therefore selected edges that were: i) present in at least 30% of subjects; and ii) were amongst the *τ* % strongest edges (based on the median streamline count) to achieve a desired connectome density. We note that multiple other thresholding approaches are available (79–81), and there is no consensus as to which works best for different datasets. Since the desired connection density is arbitrary, we examined our main results across a range of densities: *τ* = 15%, 20%, 25% for 360 region parcellation and *τ* = 5%, 10%, 15% for the higher resolution parcellation of 500 regions. We note that the actual connection density of the human connectome remains unknown, and we chose these thresholds to span a range commonly studied in the literature.

The connection matrix resulting from our thresholding procedure was then used as a binary mask for selecting edges for the heritability, gene expression analyses and generative modeling. This masking procedure thus restricted individual variability in the binary topology of connectomes across individuals (indeed, in healthy individuals such topology should be highly conserved). For heritability analysis we used this group-representative connectome as a mask to extract FA-based connection weights and also repeated the analysis using streamline count as a measure of connection strength.

### Rich-club organization

The connectivity of each region (node) in a network can be quantified by counting the number of connections to which it is attached. This measure is known as node degree. At a particular degree threshold, *k*, nodes can be labelled as hubs (degree > *k*) or nonhubs (degree ≤ *k*), subsequently classifying all connections within the network as ‘rich’ (connection between two hubs), ‘feeder’ (connection between a hub and a nonhub), and ‘peripheral’ (connection between two nonhubs) [see, Fig. 1A, (5)]. To quantify the inter-connectivity between hub regions within a binary brain connectivity network, we used the topological rich-club coefficient *Φ*(*k*):

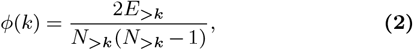

where *N*_>*k*_ is the number of nodes with degree > *k*, and *E*_>*k*_ is the number of edges between nodes with degree > *k* (82). Therefore, the rich-club coefficient quantifies the density of the subgraph comprising nodes with degree higher than the hub-defining threshold *k*.

Since nodes with higher degree make more connections, and can thus be expected to have a higher connection density compared to other nodes, we compared the *Φ*(*k*) of the empirical network to the mean value across a 1000 randomized null networks, *Φ*_rand_(*k*), generated by rewiring the edges of the empirical network while retaining the same degree sequence, using the `randmio_und` function from the Brain Connectivity Toolbox (83), rewiring each edge 50 times per null network. This randomization method is commonly used in the literature (3–5, 7). Alternative approaches that preserve both the degree sequence and connectome wiring cost (56, 84, 85) can be used to test rich-club organization in relation to geometric influences on connectome organization.

To assess whether the connections between high-degree nodes were also more likely to have stronger connection weights than expected by chance, we evaluated the weighted rich-club coefficient (86):

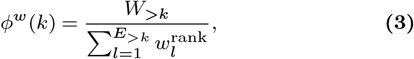

Where *W*_>*k*_ is the sum of weights in the sub-graph with degree higher than *k*, and the denominator is the total sum of *l* strongest weights in the network. As a null model for the weighted rich-club coefficient, we separate the definitions of weighted and topological rich-club coefficients by randomly reassigning weights within the network while preserving the binary topology (87) (instead of rewiring the links).

In both binary and weighted cases, we computed the normalized rich-club coefficient *ϕ*_norm_(*k*) as the ratio between the rich-club coefficient in the empirical network and the mean rich-club coefficient in the set of the corresponding randomized networks:

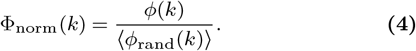

Values of Φ_norm_ > 1 indicate rich-club organization, where high-degree nodes are more densely interconnected (in a case of the topological rich-club) or have higher weights (in a case of the weighted rich-club) than be expected by chance. The statistical significance of the result is assessed by computing a *p*-value directly from the empirical null distribution of the 1000 randomized networks, *Φ*_rand_(*k*), as a permutation test (5). We note that in all our analyses, we estimated node degree using SC-weighted connectomes. Where indicated, FA-weighted connectomes were used in analyses of connectivity weights.

### Communicability

We investigated the topological centrality of rich links using a measure called communicability (88), estimated across a range of degree thresholds. The communicability, *C*_*ij*_, between a pair of nodes *i* and *j*, is calculated by accounting for all possible paths of length *l* between the nodes, weighted as 1/*l*!, so that shorter paths make a stronger contribution to the overall score. The communicability, *C*_*ij*_, for a binary matrix *A* is formally defined as:

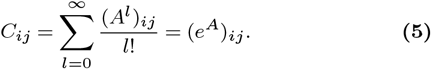

In a weighted network, communicability, 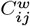, is defined using a weighted adjacency matrix *W* :

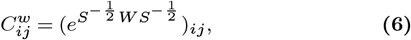

where 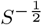 is the diagonal matrix with elements 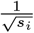 and *s*_*i*_ is the strength of node *i*. We estimated the mean binary and weighted communicability for rich links, as a function of the hub-defining threshold *k*, to evaluate whether rich links are topologically central within the human connectome (Figs. S1E,F). An advantage of communicability is that, unlike classic measures of centrality, it does not assume that information is routed exclusively along shortest paths in the network, which is likely to be an inappropriate assumption for brain networks (2, 89).

### Heritability analysis

The HCP diffusion imaging dataset includes 117 pairs of genetically confirmed monozygotic (MZ) twin pairs together with 69 of their non-twin siblings, as well as 60 dizygotic (DZ) same-sex twin pairs and 48 of their non-twin siblings. For each twin pair with more than one non-twin sibling, we selected one sibling at random (demographic details summarized in the Table 1). Only twin pairs where both twins had genetically-verified zygosity were included in the heritability analysis.

**Table 1.** Demographic data for twin groups and their non-twin siblings. MZ – monozygotic twins, DZ – dizygotic twins. Age is displayed in years: mean ± SD.

| Zygosity | Number of subjects | Sex (F/M) | Age |
| --- | --- | --- | --- |
| MZ twins | 117 pairs | 69/48 | 29.3 ± 3.3 |
| MZ non-twin siblings | 69 | 34/35 | 29.1 ± 4.2 |
| DZ twins | 60 pairs | 33/27 | 28.8 ± 3.5 |
| DZ non-twin siblings | 48 | 24/24 | 29.1 ± 4.0 |

**Table 2.**
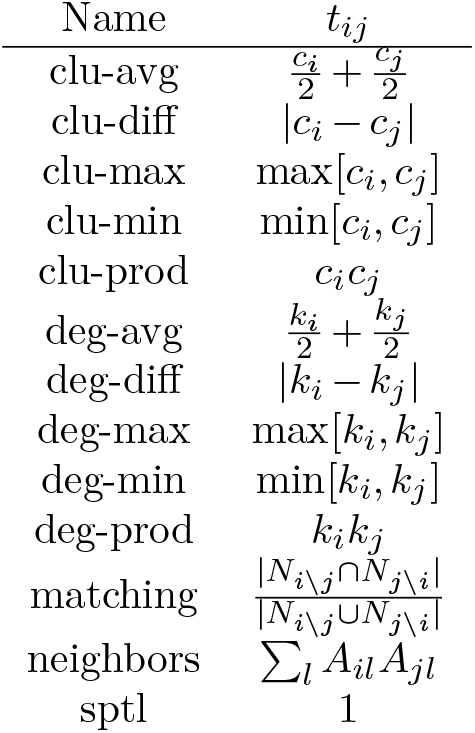
A list of topological terms, *t*_*ij*_, used in the generative models [see Eq. 8].

Heritability analysis relies on the assumption that both shared genetic factors and common environment contribute to phenotypic similarity between twins within a pair, whereas unique environmental factors and non-shared genetic effects contribute to the differences observed between them. In the classical twin design, MZ twins are assumed to be genetically identical whereas DZ twins on average share half of their DNA, which is similar to non-twin siblings. Structural equation modeling can thus be used to decompose phenotypic variance and covariance in any particular trait into additive genetic (A), common environmental (C), and unique environmental (E) influences. Considering that twins raised together might have experienced a more similar environment compared to their non-twin siblings, including a set of non-twin siblings into the analysis allows us to separate the common environmental contributions into twin-specific (T) and twin non-specific (C) common environmental factors.

We used the binary group-representative cortical connectome mask described above to extract FA-weighted edges and applied standard structural equation modeling (SEM) to every connection in the connectome using OpenMx software (90, 91) in R. The analysis reported in the main text was performed on the 360 region (37) cortical connectome at 20% density (12 924 unique connections) using FA as a connection weight. The analyses were subsequently reproduced using SC (Fig. S2C,D) and at different connectome densities (Fig. S3A-C) and using a higher resolution 500-region random cortical parcellation at 5%, 10% and 15% densities (Fig. S3D–F).

A range of biometric models – ACTE, ACE, AE, CE, E – were fitted to each edge defined by the group connectome mask in order to find connection-specific maximum likelihood estimates of additive genetic (A), twin-specific common environmental (T), twin non-specific common environmental (C) and unique environment (E) factors, using age and sex as covariates. Outlying connection weights for each analysis were removed using the boxplot function in R by keeping data points (*w*) in a range *Q*1 − 1.5 × *IQR* < *w* < *Q*3 + 1.5 × *IQR* where *Q*1 and *Q*3 are the first and third quartiles respectively and *IQR* is the interquartile range. The Akaike information criterion (AIC) (92) was used to compare the goodness of fit of all tested models in order to find the most parsimonious model. For each edge, the model with the lowest AIC was selected. Consequently, the narrow-sense heritability (the proportion of variance attributable to additive genetic factors, referred to as heritability) was estimated for each connection using the best-fitting model. We also show heritability results using parameter estimates from the full ACTE model to ensure that our findings cannot be explained by our model selection procedure (Fig. S2B) and verify that outlier exclusion did not affect our findings (Fig. S4D-F).

### Gene expression data

We used brain-wide gene expression data from the Allen Human Brain Atlas (AHBA), which consists of microarray expression measures in 3702 spatially distinct tissue samples taken from six neurotypical postmortem adult brains (40). Different brain regions were sampled across each of the six AHBA donors to maximise spatial coverage, resulting in approximately 400–500 tissue samples in each brain. The samples were distributed across cortical, subcortical, brainstem and cerebellar regions, measuring the expression levels of 58 692 probes quantifying the transcriptional activity of 20 737 genes. Considering that only two out of six brains were sampled from both left and right hemispheres whereas the other four brains had samples collected only from the left hemisphere, we focused our analyses on the left cortex only.

The pre-processing procedures applied to the data are outlined below and the choices detailed in (35). Briefly, probe-to-gene annotations were first updated using the Re-Annotator toolbox (93) resulting in the selection of 45 821 probes corresponding to the total of 20 232 genes. Second, tissue samples annotated to the brainstem and cerebellum were removed. Then, intensity-based filtering (35) was applied in order to exclude probes that do not exceed background noise in more than 50% of samples, excluding 13 844 probes corresponding to 4486 unique genes. Afterwards, a representative probe for each gene was selected based on the highest correlation to RNA sequencing data in two of the six brains (94). Gene expression samples were assigned to regions-of-interest by generating donor-specific grey matter parcellations and assigning samples located within 2 mm of the parcellation voxels. To increase the accuracy of assigning samples to regions, the samples were first divided into four separate groups based on their location: hemisphere (left/right) and structure assignment (cortex/subcortex), so samples listed as coming from the left cortical hemisphere in the AHBA ontology are only mapped to left cortical voxels of the parcellation (applying a 2 mm distance threshold, almost 90% of all cortical and subcortical samples were assigned to a non-zero voxel of the parcellation). Then, samples assigned to the subcortical regions as well as the right hemisphere were removed. Finally, gene-expression measures within a given brain were normalized first by applying a scaled robust sigmoid normalization for every sample across genes and then for every gene across samples in order to evaluate the relative expression of each gene across regions, while controlling for donor-specific differences in gene expression [see (35) for a validation]. Normalized expression measures in samples assigned to the same region were averaged within each donor brain and aggregated into a region by gene × matrix consisting of expression measures for 10 027 genes over 180 (left hemisphere, HCP parcellation) and 250 regions (left hemisphere of the random parcellation), respectively.

Distances between region pairs that were subsequently used to account for the spatial effects on transcriptional coupling were estimated on the cortical surface (pial) using the annotation files for each parcellation mapped onto the spherical representation of the `fsaverage` cortical surface. First, all the vertices that correspond to a particular region of interest in the spherical representation were identified and their centroid coordinates were calculated. Then the centroid coordinates were mapped to the fsaverage cortical surface and the distances between each pair of regions were calculated using the toolbox `fast_marching_toolbox` in MATLAB.

### Transcriptional coupling

The result of the above mapping of AHBA data was an expression profile for each brain region, quantifying transcriptional activity across 10 027 genes. We used these profiles to quantify transcriptional coupling, or correlated gene expression (CGE), between every pair of regions. We defined CGE as the Pearson correlation between the normalized expression measures of the genes available after pre-processing (*n* = 10027). As shown in Fig. S5A and described in (35), CGE exhibits a strong spatial autocorrelation that can be approximated as an exponential relationship with separation distance,, such that regions located in close proximity to each other share more similar gene expression. To investigate whether CGE differs between different topological classes of connections beyond any low-order spatial effect, we need to ensure that the distance between regions alone is not informative of their CGE. Otherwise, any similarity between region pairs will be driven by a mixture of two factors: the CGE signature that is reflective of the spatial gradient and CGE signature that corresponds to the edge-specific properties. To account for the low-order spatial effect we fitted an exponential function with form *r*(*d*) = *Ae*^−*d/n*^ + *B*. The parameters *A* = 0.64, *B* = −0.19 and *n* = 90.4 capture the trend well, allowing us to retain the residuals for further analysis (Fig. S5B), defined as 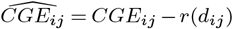. These distance-corrected residual CGE values were used in all CGE analyses.

To evaluate transcriptional coupling for different connection types, for every edge within the connectivity matrix, we assigned a distance-corrected CGE measure. At each degree threshold, *k*, for defining hubs (nodes with degree> *k*), we then computed the average CGE of rich, feeder, and peripheral links. Significant increases in the CGE for a given link type compared to the rest of the network were evaluated using a one-sided Welch’s *t*-test (*p* < 0.05).

### Gene contribution score

To determine which functional gene groups contribute the most to any observed differences in CGE across different link types in the brain, we quantified the degree to which each gene contributes to the overall CGE between a pair of regions, following prior work (7):

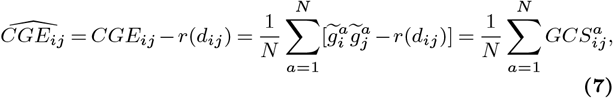

where *N* is the number of genes 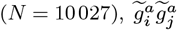 the product of the *z*-score normalized expression values for gene *a* in regions *i* and *j*, and *r*(*d*_*ij*_) is the previously defined spatial autocorrelation effect approximated as an exponential line (Fig. S5). Therefore, the gene contribution score between a pair of regions *i* and *j* for gene *a* was defined as 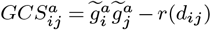

We then assigned each gene a *t*-statistic quantifying the increase in GCS for rich compared to peripheral links (GCS_t stat_), as these two groups constitute the most distinct link types. A high value indicates increased CGE in rich compared to peripheral links. These *t*-statistic measures were used in the enrichment analyses as gene scores for determining whether any functional gene groups made a stronger contribution to CGE than others.

### Cell-specific genes

Given that the AHBA assays gene expression using bulk tissue samples, it is possible that regional variations in cellular architecture drive differences in CGE between different link types. To test this hypothesis, we conducted a second CGE analysis focused on subsets of genes that have previously been identified as cell-specific markers. The set of cell-specific genes was compiled based on data from five different single-cell studies that used postmortem cortical samples of human postnatal subjects. Genes identified in each study as a cell-specific marker or as specifically enriched within a cell type were aggregated into study-specific lists (44–47). In the case of (43) where the normalized gene expression values were available for each cell type, we identified enriched genes as those with an average Fragments per kilobase million, FPKM> 5 and at least a four-fold enrichment over other cell types, as per authors recommendations. We then assigned genes within each of the resulting study-specific gene lists to one of seven canonical cell classes: astroglia, endothelial cells, excitatory neurons, inhibitory neurons, oligodendrocytes, and oligodendrocyte progenitor cell restricting each cell-class list to only contain genes unique to that class.

### Gene-set enrichment analysis using gene score resampling

Gene-set enrichment analyses assess whether any functionally related groups of genes, annotated using Gene Ontology (GO), are associated with a selected phenotype. Every gene in our sample (*n* = 10027 genes) was assigned a *t*-statistic score quantifying its contribution towards the increase in GCS for rich links relative to peripheral (GCS_t−stat_). Using these scores, we determined which specific functional groups of genes contribute to the observed increase in correlated gene expression. Functional gene group analysis was performed using version 3.1.2 of ErmineJ software (95). Gene ontology (96) annotations were obtained from GEMMA (97) as Generic_human_ensemblIds_noParents.an.txt on December 9, 2019. Gene Ontology terms and definitions were acquired from the archive.geneontology.org/latest-termdb/go_ daily-termdb.rdf-xml.gz on January 13 2020. We performed gene score resampling (GSR) analysis on the *GCS*_t−stat_ scores testing the biological process GO categories with 5 to 100 genes available using the mean *t*-statistic score across genes to summarize the GO category and applying full resampling with 10^6^ iterations. The resulting *p*-values were corrected across 6201 GO categories, controlling the false discovery rate (FDR) at 0.05 using the method of Benjamini and Hochberg (98).

Recent work indicates that the default null models used in such analyses are insufficiently constrained for spatially embedded transcriptomic atlas data (50). This problem can lead to inflated significance for some GO categories when testing for spatial correlations between regional variations in gene expression patterns and measures of brain structure or function. The extent to which this problem generalizes to phenotypes defined for pairs of regions, such as the connectivity metrics considered here, is unclear. We nonetheless suggest caution in interpreting the findings of this analysis, as appropriate null models for the analysis of pairwise phenotypes have not yet been developed. We report the enrichment findings to test for consistency with prior findings in the mouse (7).

### Microstructural profiles

Our CGE analysis of cell-specific genes indicated that connected hubs have more similar cellular composition than other region pairs. To independently verify this result, we estimated the microstructural profile covariance (MPC) between each pair of regions using the BigBrain atlas, which is a Merker-stained 3D volumetric histological reconstruction of a human brain (48, 49). MPC was estimated using methods described in [(49), see https://github.com/MICA-MNI/micaopen/tree/master/MPC]. In brief, the MPC procedure involved constructing 16 equivolumetric surfaces between the pial and white matter boundaries, followed by systematic sampling of the intensity values along these surfaces at 163 842 matched vertices per hemisphere. The intensity profiles, reflecting depth-wise changes in cellular density and soma size, were corrected for the midsurface *y*-coordinate to account for an anterior–posterior increase in intensity values across the BigBrain related to coronal slicing and reconstruction. Standardized residual intensity profiles were averaged within areas of the HCPMMP1 (*n* = 360) (37) and random (*n* = 500) parcellations. We quantified cytoarchitectural similarity between cortical areas by correlating areal intensity profiles (covarying for cortex-wide mean intensity profile), thresholding to retain only positive values (*r* > 0) and applying a log transformation, resulting in the measure of microstructural profile covariance (MPC) (49). We repeated the same analysis using the 500-region random parcellation.

Notably, this analysis did not replicate the elevated MPC for rich links seen with the HCPMMP1 atlas (compare Fig. 3H with Fig. S8B). This discrepancy likely reflects the fact that the HCP-MMP1 parcellation more closely approximates boundaries between functional zones of the cortex, as it is based on a fusion of multimodal imaging data (37). The random parcellation makes no attempt to capture such boundaries and may blur different cytoarchitectonic regions within the same network node, thus resulting in noisier MPC estimates. In this way, the MPC results appear to depend on accurate approximation of cytoarchitectonic boundaries in cortex.

### Models of brain network wiring

To evaluate the role of stochastic processes in shaping connectome architecture, we evaluated a series of generative models of network wiring that have the general form:

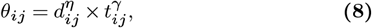

where *θ*_*ij*_ is a score that weights the probability of connecting nodes *i* and *j, d*_*ij*_ is the Euclidean distance between nodes *i* and *j*, and *t*_*ij*_ is some topological property of nodes *i* and *j* or an edge between them. This topological term modulates the probabilities of forming an edge along with wiring cost (operationalized as *d*_*ij*_). Numerous topological properties have been evaluated for *t*_*ij*_ in past work (32, 33), and we consider these same models here. A summary is provided in Table 2.

*c* is the local clustering coefficient, *k* represents node degree, *A*-adjacency matrix and *Ni*_*\*_*j* - neighbors of the node *i* excluding node *j*. The exponents, *η* and *γ*, act as weights on the distance and topological terms, respectively (32, 33). At each iteration, the computed connection score, *θ*_*ij*_, is used to calculate the probability of a given edge, (*i, j*), being formed in that iteration, as *P*_*ij*_ = θ_*ij*_ /Θ, where Θ is the sum of *θ*_*ij*_ over all edges that have not yet been formed. Thus, at a given iteration, the model calculates the probability of each edge forming based on its distance and the current value of its topological term, *t*_*ij*_. This topological value is recalculated at each iteration. Edges are added iteratively until the total number of edges is equal to the number of edges in the empirical connectome. Due to computational burden, and in line with prior analyses (33), we fitted models to a single (left) hemisphere connectome defined using the HCPMMP1 (37) parcellation, containing 5025 unique edges (20% whole cortex connectome density).

As per prior work (33), we quantified model performance using the Kolmogorov-Smirnov (KS) statistic. The KS statistic quantifies the distances between distributions of key topological statistics of the network; as such, lower values indicate better model fit. We focused on four key metrics: node-level distributions of degree, clustering, and betweenness, and the edge-level distribution of connection distance (33). The quality of model fit was defined as the maximum KS value across all four distributions; that is, model performance is defined by the property that is fitted most poorly. In principle, any number of other topological parameters could be used in this objective function, but these are some of the most widely used to characterize brain networks and were employed in prior work evaluating the same models (32, 33)

We optimize the free parameters *η* and *γ* as per previous work (33). Specifically, we randomly sample the parameter space and evaluate the fits of the resulting networks. We consider *η* values in the range from −4 to 4, allowing both positive and negative contributions for the topological terms, whereas wiring cost is always penalized with *γ* values ranging from −15 to 0. After sampling 2000 points in this space, Voronoi tessellation is then used to identify areas – or cells – of this space where the parameters produce networks with the best fits, as defined by the KS statistic. We then preferentially sample from cells with better fits. This procedure is repeated four times so that the algorithm gradually converges on an optimum. We ran each generative model on the group connectome 10 000 times and then evaluated each different model by comparing the 100 lowest energy values obtained from the optimization procedure.

For our analysis, we draw a critical distinction between the distribution and sequence of a topological property. The distribution refers to how a property is statistically distributed across the nodes of the network. The sequence refers to the exact assignment of a particular value to individual nodes or edges; in other words, how the property is spatially embedded in the brain. It is possible that two networks may have similar distributions for a given property, but very different underlying sequences.

The models we consider here are optimized to match distributions, not sequences. As we are specifically interested in understanding the mechanisms driving the precise way in which hubs are connected, and given evidence that the specific anatomical location of network hubs has important implications for network dynamics (56), we seek to evaluate whether the models can not only generate networks with hubs, which would be shown by accurate fitting of the degree distribution, but also whether they yield hubs in the same anatomical regions as the empirical data, which would be shown by accurate fitting of the degree sequence. To this end, we additionally evaluate the correlation between the degree sequences of the empirical and synthetic networks using the Spearman correlation coefficient, *ρ*. A high correlation between the model and data implies that hubs are located in the same anatomical regions across the two networks. Conversely, a low correlation indicates that the model does not accurately capture the spatial embedding of connectivity in the connectome. Put simply, a low correlation implies that the hubs in the model network reside in anatomical locations that differ from the actual connectome.

An important consideration is that the correlation between model and empirical degree sequences was not a part of the objective function used in model fitting. We fitted the models using topological distributions and then evaluated their performance in capturing the empirical degree sequence. This procedure allows us to examine how well these models, as traditionally implemented, capture spatial properties of hub connectivity. However, this procedure also raises the question of whether it is possible to obtain a higher degree sequence correlation if model parameters are chosen to optimize this specific quantity. We therefore repeated the analysis after replacing the objective function with one that maximized the similarity between model and empirical degree sequences. Specifically, we optimized the Spearman correlation between model and empirical degree sequences with no other constraints to give the models the best possible chance of reproducing the empirically observed spatial topography of hub regions. The results of this analysis are shown in Fig. S11. Qualitatively similar results were obtained when optimizing the Pearson correlation between model and empirical degree sequences.

Across the 13 models evaluated in our analysis, we find that the best-fitting model is the ‘deg-avg’ model, which involves a tradeoff between forming connections between highly connected nodes (i.e., node pairs with high average degree) and penalizing long-range connections (i.e., minimizing wiring cost). This result differs from past work, in which a homophilic attachment trade-off model that balances wiring cost with a preference for forming connections between nodes with similar neighbors offered the best fit to empirical connectome data (32, 33). This discrepancy may be related to our use of a higher resolution network parcellation, a connectome mapped at a different connection density, using a different tractography algorithm, and/or a different diffusion MRI processing pipeline. Investigating the effect of these factors on modeling results is an important direction of future work, but these potential effects do not change the substantive point of our results that current stochastic models do a poor job of reproducing the spatial embedding of hub connectivity, as across 260 000 runs of the 13 different models considered, no degree sequence correlation exceeded *ρ* = 0.3. We also note that our model fits (Figs 4B–E) are in the same range as those reported by Betzel et al. (33), indicating that our discrepant results are not due to differences in model accuracy.

Models incorporating transcriptional information are constructed using the same general form (Eq.8) while replacing one of the terms with pairwise distance-corrected CGE values *g*_ij_, weighted using a parameter *λ* varying in the range [0,200] that favors forming connections between regions with higher CGE.

While our analysis mimics prior comprehensive evaluations of generative network models based on cost-value trade-offs (32, 33), other formulations and approaches are also possible (99–101). It is also possible to define 3-parameter or higher-order models that incorporate spatial, topological and genetic constraints, but we have focused on the simpler 2-parameter form here to enable efficient optimization and fair model comparison. It is also possible that alternative, non-genetic wiring rules may yield improved model performance, and thus we cannot completely rule out a role for stochastic processes in shaping hub connectivity. Indeed, more abstract models suggest that stochastic wiring, acting in concert with developmental changes in brain geometry and heterogeneous timing of connection formation across regions, can indeed generate networks with brain-like properties (34, 59, 102). However, a framework for directly fitting such models to human DWI data has not yet been developed.

## ACKNOWLEDGEMENTS

We would like to thank Richard Betzel for sharing the optimization code for the generative modeling. This work was supported by the MASSIVE HPC facility (www.massive.org.au).

