## Supplementary file for "Genetic influences on hub connectivity of the human connectome"

*Contributed equally to this work
[1]The Turner Institute for Brain and Mental Health, School of Psychological Sciences, and Monash Biomedical Imaging, Monash University, Victoria, Australia.
[2]School of Physics, The University of Sydney, Camperdown, NSW 2006, Australia.
[3]Translational Neurogenomics Laboratory, QIMR Berghofer Medical Research Institute, Brisbane, Queensland, Australia.
[4]McConnell Brain Imaging Centre, Montreal Neurological Institute and Hospital, McGill University, Montreal, Canada.
[5]Murdoch Children's Research Institute, Melbourne, Victoria, Australia.
[6]Department of Human Genetics, Radboud University Medical Center, Donders Institute for Brain, Cognition and Behaviour, Nijmegen, The Netherlands
[7]Department of Psychiatry, University of California San Diego, La Jolla, CA, USA
[8]Center for Brain and Cognition, Computational Neuroscience Group, Department of Information and Communication Technologies
[9]Universitat Pompeu Fabra, Roc Boronat 138, Barcelona, 08018, Spain
[10]Institució Catalana de la Recerca i Estudis Avançats (ICREA), Passeig Lluís Companys 23, Barcelona, 08010, Spain
[11]Department of Neuropsychology, Max Planck Institute for Human Cognitive and Brain Sciences, 04103 Leipzig, Germany
[12]Department of Psychiatry, Radboud University Medical Center, Donders Institute for Brain, Cognition and Behaviour, Nijmegen, The Netherlands

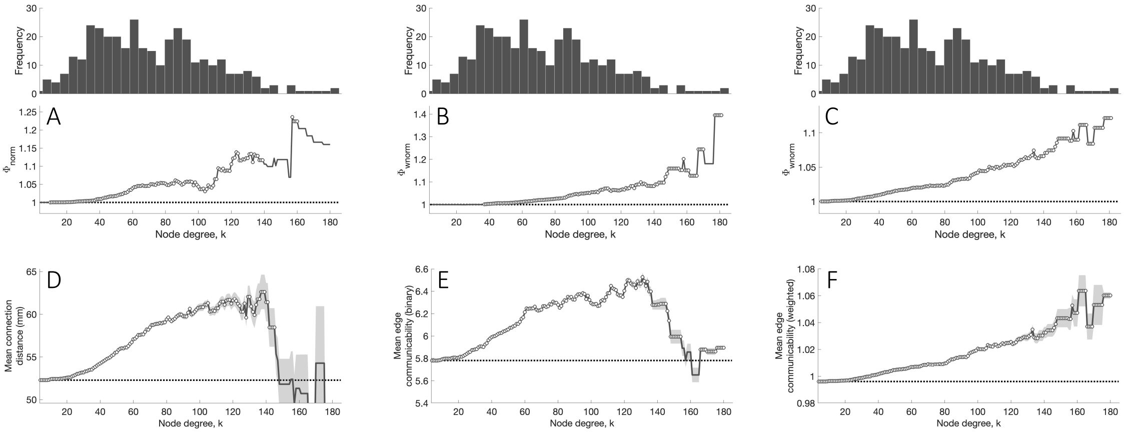


Figure S1: **Rich-club organization of the connectome**. Top: the degree distribution across regions displays a long-tail indicating the presence of a relatively small number of highly connected regions - brain network hubs. Bottom: **(A)** Normalized topological rich-club coefficient $\phi_{\mathrm{norm}}$; $\phi_{\mathrm{norm}} > 1$ indicates that hubs are more densely interconnected than expected by chance, which is consistent with rich-club organization. Connections between nodes in the tail of the degree distribution, particularly spanning $105 < k < 146$, demonstrate consistently increasing $\phi_{\mathrm{norm}}$. This threshold range corresponds to the topological rich-club regime; circles indicate values that are significantly higher compared to an ensemble of 1000 degree-matched null networks ($p < 0.05$). **(B)** Normalized weighted rich-club coefficient using log-transformed streamline counts as connection weights. **(C)** Normalized weighted rich-club coefficient using tract-averaged fractional anisotropy as connection weights. $\phi_{\mathrm{wnorm}} > 1$ indicates that connections between hubs are stronger than expected by chance; circles indicate values that are significantly higher than an ensemble of 1000 topology-matched null networks ($p < 0.05$). **(D)** Mean Euclidean distance between connected hub regions as a function of the degree, $k$, at which hubs are defined (as regions with degree $> k$). The mean connection distance across all network links is shown as a dotted black line; circles indicate values significantly greater than all other pairs of connected regions (right-tailed Welch's $t$-test, $p < 0.05$). Mean normalized binary **(E)** and weighted **(F)** connection communicability as a function of the node degree, $k$. Normalized edge communicability is calculated as the ratio between connection communicability in the empirical connectome and an ensemble of 1000 degree-matched null networks. Shaded areas denote standard error of the mean across edges at a selected degree threshold. The mean normalized edge communicability across all network links shown as a dotted black line; circles indicate values significantly greater than across all other pairs of connected regions (right-tailed Welch's $t$-test, $p < 0.05$). Measures of wiring cost (distance) and centrality all increase as a function $k$, indicating that connections between hubs are among the most costly and topologically central in the brain.

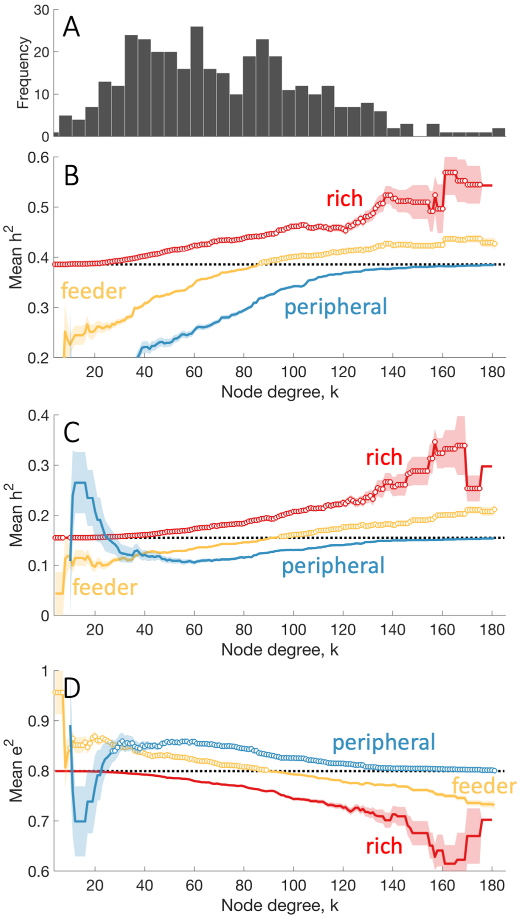


Figure S2: **Mean heritability estimates derived from the full ACTE model**. **(A)** The degree distribution of the representative group-level connectome. **(B)** Mean edge heritability for rich (hub-hub), feeder (hub-nonhub), peripheral (nonhub-nonhub) connections as a function of degree threshold, $k$ used to define hubs. **Genetic and unique environmental influences on connectivity strength estimated using streamline count for each connection type**. Mean heritability **(C)**, and unique environmental factors **(D)** for rich (hub-hub), feeder (hub-nonhub), peripheral (nonhub-nonhub) connections as a function of degree threshold, $k$ used to define hubs. In each panel the mean of the measure across all network links is shown as a dotted black line. Shaded area corresponds to the standard error of the mean. Circles indicate a statistically significant increase in the measure in a given link type compared to the rest of the network (one-sided Welch's $t$-test, uncorrected $p < 0.05$).

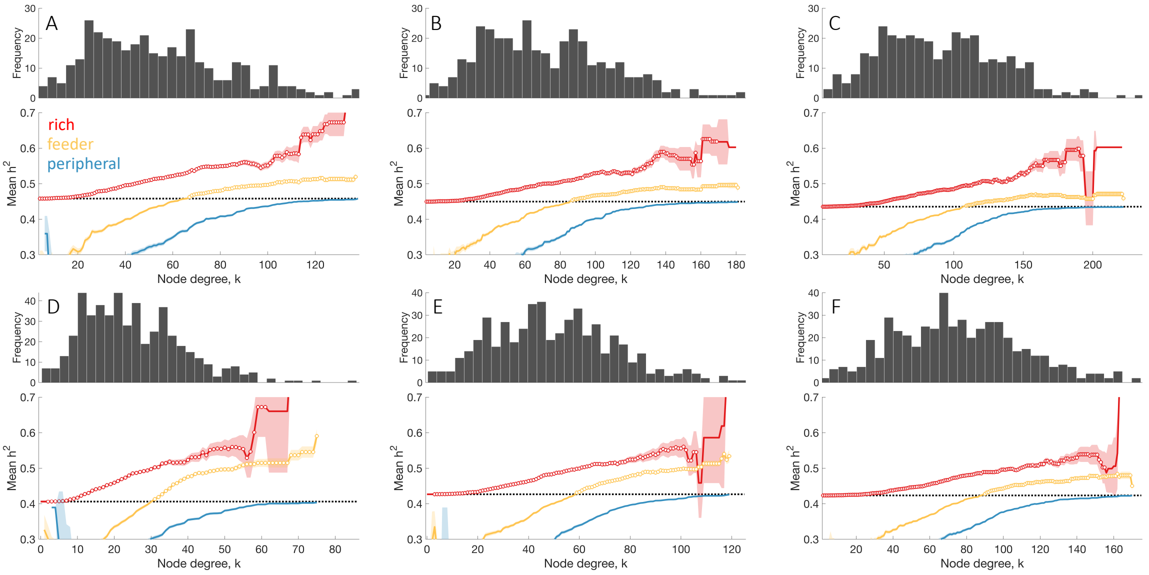


Figure S3: **Rich links show consistently higher heritability across different connectome processing options.** Top row shows results for the 360-region HCPMMP1 parcellation at 15% **(A)**, 20% **(B)**, 25% **(C)** connectome density; bottom row shows results for the random 500-region parcellation at 5% **(D)**, 10% **(E)**, 15% **(F)** connectome density. In each plot top: the degree distribution of the group-level connectome. Bottom: mean edge heritability for rich (hub-hub), feeder (hub-nonhub), peripheral (nonhub-nonhub) connections as a function of degree threshold, $k$ used to define hubs. The mean heritability across all network links is shown as a dotted black line. Shaded area corresponds to the standard error of the mean, circles indicate a statistically significant increase in heritability in a given link type compared to the rest of the network (one-sided Welch's $t$-test, uncorrected $p < 0.05$).

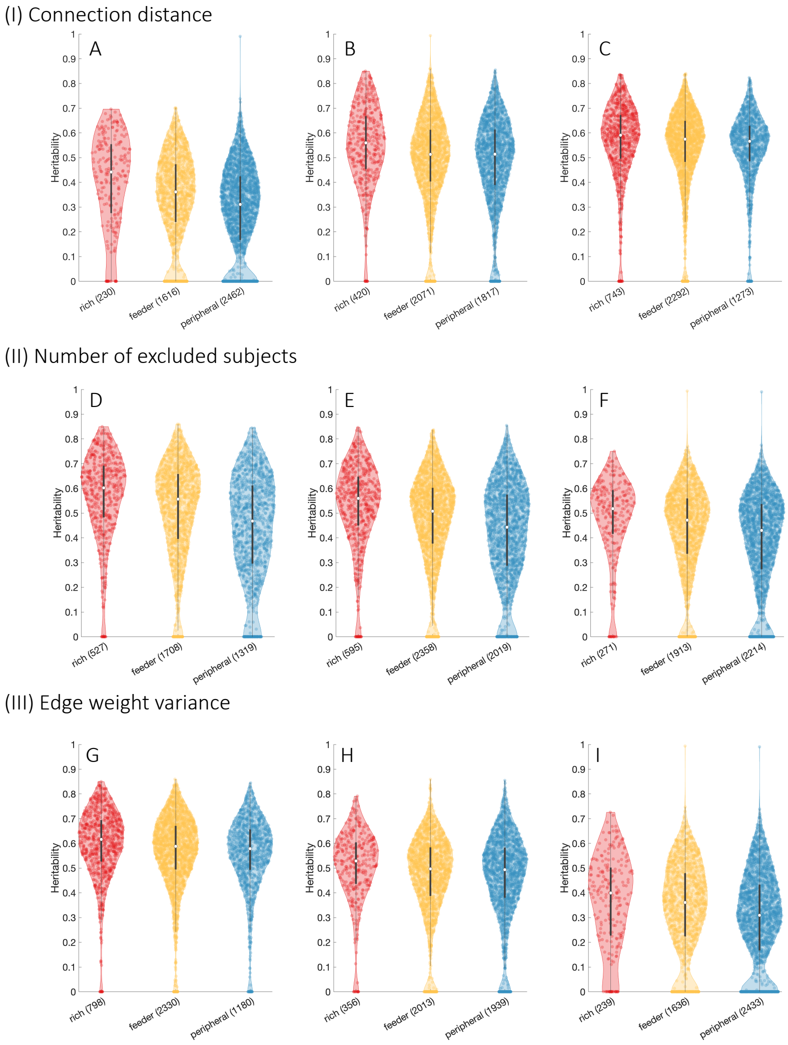


Figure S4: **(I) Heritability across different connection lengths.** To examine whether preferential genetic influences on rich links can be explained by connection distance, we plot distributions of edge-specific heritability estimates within three connection distance bins [short ($< 76mm$, **(A)**), medium ($76mm - 114mm$, **(B)**) and long ($> 114mm$, **(C)**). Distances are estimated as the average tract length for rich, feeder and peripheral links]. Rich links show significantly higher heritability compared to both feeder and peripheral links across all three distance ranges (one-sided Welch's $t$-test, all $p < 1.9 \times 10^{-3}$), suggesting that distance effects cannot explain our findings. **(II) Heritability across groups of edges with a different number of outlier exclusions.** Outlying values were excluded from heritability analyses at each edge. To examine whether the preferential genetic influence on rich links could be explained by this exclusion procedure, we group edges into three types, based on the number of excluded outlying values [low (data from $< 6$ participants excluded, **(D)**), medium (data from $6 - 14$ participants excluded, **(E)**) and high (data from $> 14$ participants excluded, **(F)**)], and plot distributions of edge-specific heritability estimates for rich, feeder, and peripheral links. Rich links show significantly higher heritability compared to both feeder and peripheral links across all three groups of edges (one-sided Welch's $t$-test, all $p < 3.1 \times 10^{-5}$). **(III) Heritability across groups of edges with different levels of phenotypic variance in connectivity strength.** To examine whether differences in the phenotypic variance of connectivity strength estimates could explain the preferential genetic influence for rich links, we plot distributions of edge-specific heritability estimates within three connection variance bins [low ($< 5.5 \times 10^{-4}$, **(G)**), medium ($< 5.5 \times 10^{-4} - 7.3 \times 10^{-4}$, **(H)**) and high ($> 7.3 \times 10^{-4}$, **(I)**)]. Rich links show significantly higher heritability compared to both feeder and peripheral links within low and medium variance ranges (one-sided Welch's $t$-test, both $p < 4.9 \times 10^{-6}$) and slightly higher heritability within the high variance range (one-sided Welch's $t$-test, $p = 0.03$).

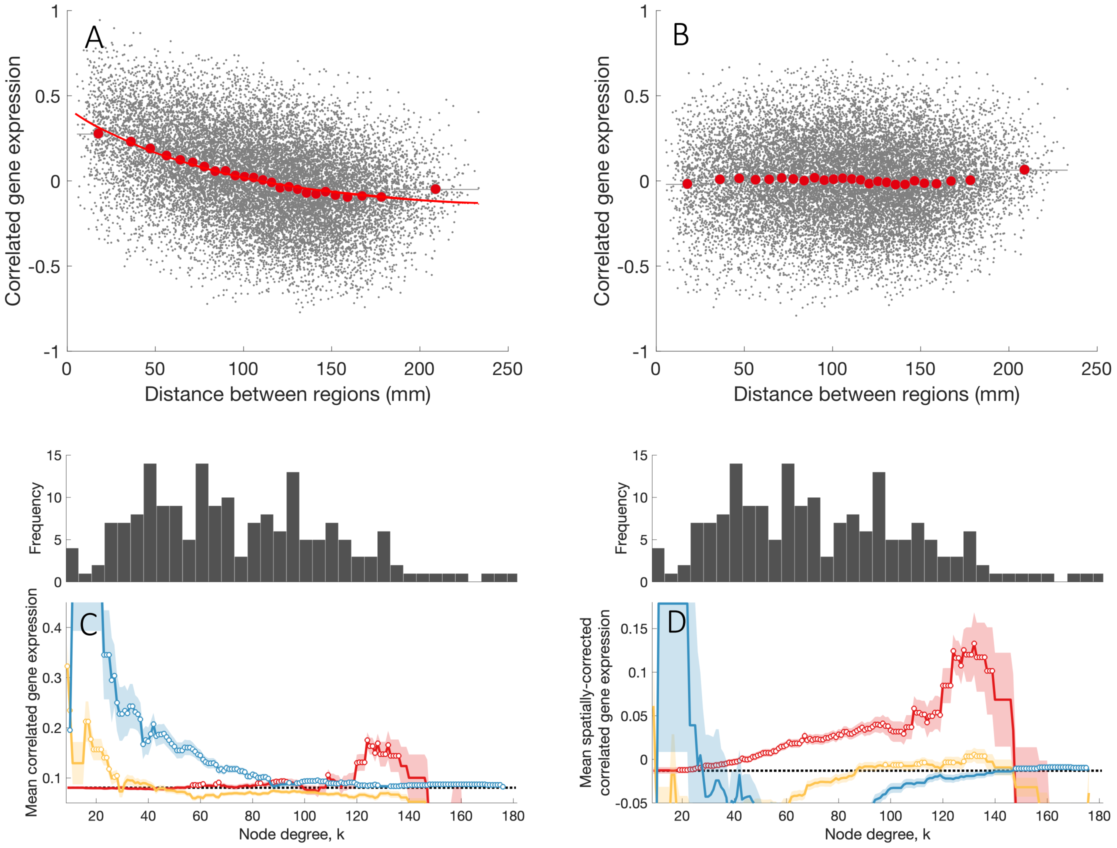


Figure S5: **Correlated gene expression demonstrates a strong spatial autocorrelation.** Relationship between correlated gene expression (CGE) and regional separation distance, estimated as geodesic distance on the cortical surface (d). **(A)** CGE as a function of the regional separation distance on the cortical surface. The red line represents an exponential fit, $r(d) = 0.64e^{-d/90.4} - 0.19$. **(B)** CGE residuals after removing the exponential trend. CGE between pairs of regions are represented in grey dots and red dots represent the mean value in 25 equiprobable distance bins. Transcriptional coupling across link types before **(C)** and after **(D)** CGE distance correction. Before distance correction peripheral links show elevated CGE which is consistent with high CGE for nearby regions. There is a CGE elevation for hub-hub pairs at high $k$, although this is attenuated compared to the corrected data. In each plot top: the degree distribution of the representative group-level connectome of brain regions in the left cortical hemisphere. Degree is computed from whole-brain connectivity. Bottom: mean CGE for rich (hub-hub), feeder (hub-nonhub), peripheral (nonhub-nonhub) connections as a function of degree threshold, $k$ used to define hubs. The mean CGE across all network links is shown as a dotted black line. Shaded area corresponds to the standard error of the mean, circles indicate a statistically significant increase in CGE in a given link type compared to the rest of the network (one-sided Welch's $t$-test, uncorrected $p < 0.05$).

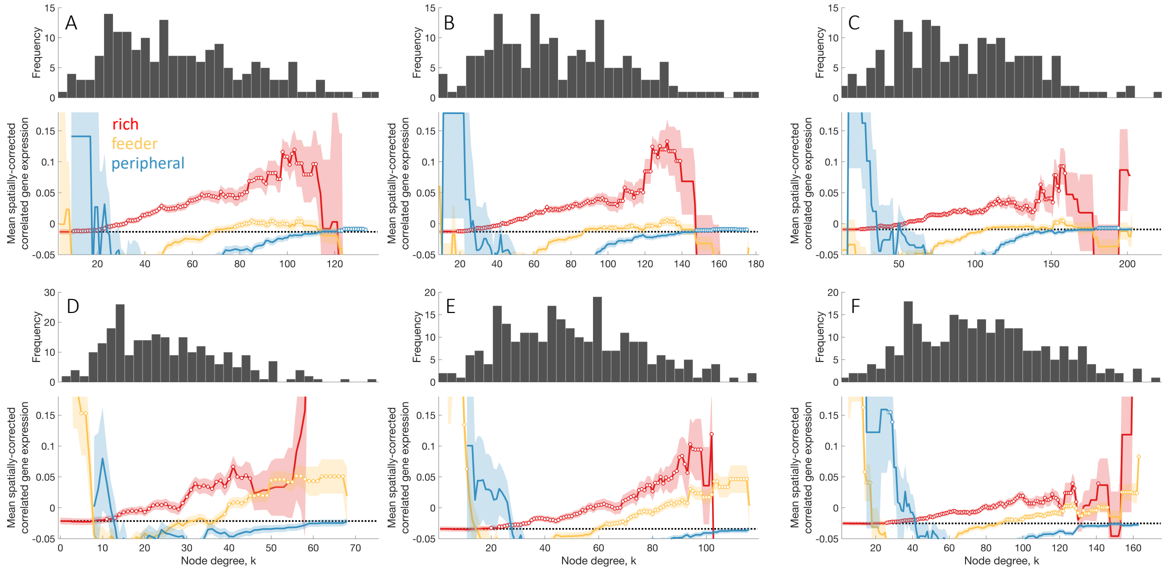


Figure S6: **Rich links show consistently higher transcriptional coupling across different connectome processing options.** Transcriptional coupling across link types for different connectome processing options. Top row shows results for the 360-region HCPMMP1 parcellation at 15% **(A)**, 20% **(B)**, 25% **(C)** connectome density. Bottom row shows results for the 500-region random parcellation at 5% **(D)**, 10% **(E)**, 15% **(F)** connectome density. In each plot top: the degree distribution of the representative group-level connectome of brain regions in the left cortical hemisphere. Degree is computed from whole-brain connectivity. Bottom: mean CGE for rich (hub-hub), feeder (hub-nonhub), peripheral (nonhub-nonhub) connections as a function of degree threshold, $k$ used to define hubs. The mean CGE across all network links is shown as a dotted black line. Shaded area corresponds to the standard error of the mean, circles indicate a statistically significant increase in CGE in a given link type compared to the rest of the network (one-sided Welch's $t$-test, uncorrected $p < 0.05$).

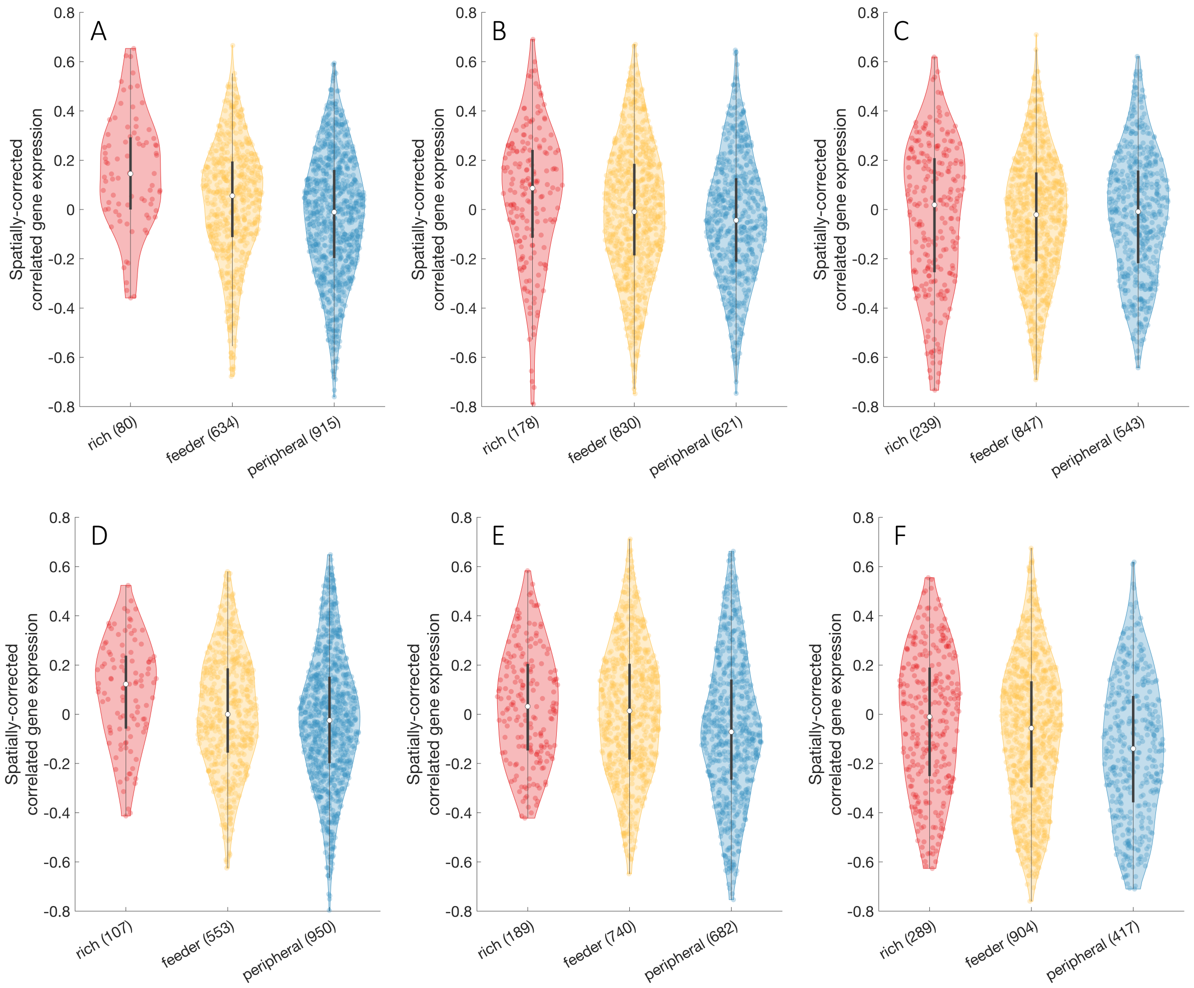


Figure S7: **Correlated gene expression across different connection lengths.** Top: CGE distributions within three distance bins [short ($< 56mm$, **(A)**), medium ($56mm - 102mm$, **(B)**) and long ($> 102mm$, **(C)**) as defined using HCPMMP1 parcellation] for rich, feeder and peripheral links. Rich links show significantly higher CGE compared to both feeder and peripheral links across short and medium distances (right-tailed Welch's $t$-test, all $p < 0.004$). There are no significant differences between CGE across link types when regions are separated by long ($> 102mm$) distances. Bottom: CGE distributions within three distance bins [short ($< 39mm$, **(D)**), medium ($39mm - 81mm$, **(E)**) and long ($> 81mm$, **(F)**) as defined using random 500 region parcellation] for rich, feeder and peripheral links. Rich links show significantly higher CGE compared to both feeder and peripheral links across short and long distances (right-tailed Welch's $t$-test, all $p < 0.006$). In the mid-range, rich and feeder links show increased CGE compared to peripheral links (right-tailed Welch's $t$-test, all $p < 5 \times 10^{-7}$). Distances are estimated as the geodesic distance on the cortical surface.

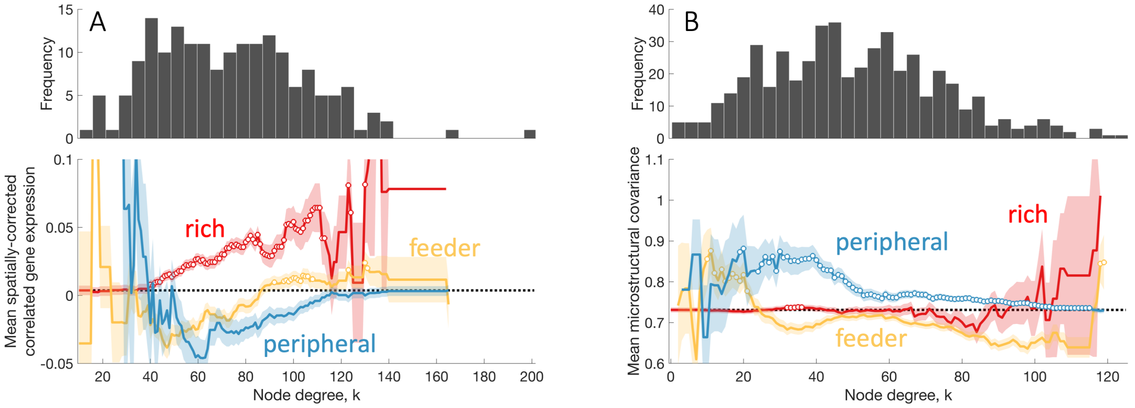


Figure S8: **Transcriptional coupling is elevated for connected brain network hubs**. Transcriptional coupling results reproduced using the Monash dataset with the HCPMMP1 parcellation and 20% connectome density. **(A)** The degree distribution of the representative group-level connectome (top). Mean CGE for rich (hub-hub), feeder (hub-nonhub), peripheral (nonhub-nonhub) connections as a function of degree threshold, $k$ used to define hubs. The mean CGE across all network links shown as a dotted black line. Shaded area corresponds to the standard error of the mean, circles indicate a statistically significant increase in transcriptional coupling in a given link type compared to the rest of the network (one-sided Welch's $t$-test, $p < 0.05$). **Rich links do not show increased microstructural similarity when using a random cortical parcellation.** **(B)** The degree distribution of the representative group-level connectome using 500 region parcellation at 10% connectome density (top). Mean microstructural profile covariance (MPC) for rich (hub-hub), feeder (hub-nonhub), peripheral (nonhub-nonhub) connections as a function of degree threshold, $k$ used to define hubs. The MPC across all network links shown as a dotted black line. Circles indicate a statistically significant increase in MPC in a given link type compared to the rest of the network (one-sided Welch's $t$-test, $p < 0.05$). The discrepancy observed between the random and HCPMMP1 parcellations (Fig. 3H) likely reflects the fact that regional borders in the latter are more closely aligned with cytoarchitectonic boundaries, as the parcellation explicitly tries to delineate functional zones of cortex based on multimodal neuroimaging data [1]. The random parcellation does not try to approximate such boundaries, resulting in noisier MPC estimates.

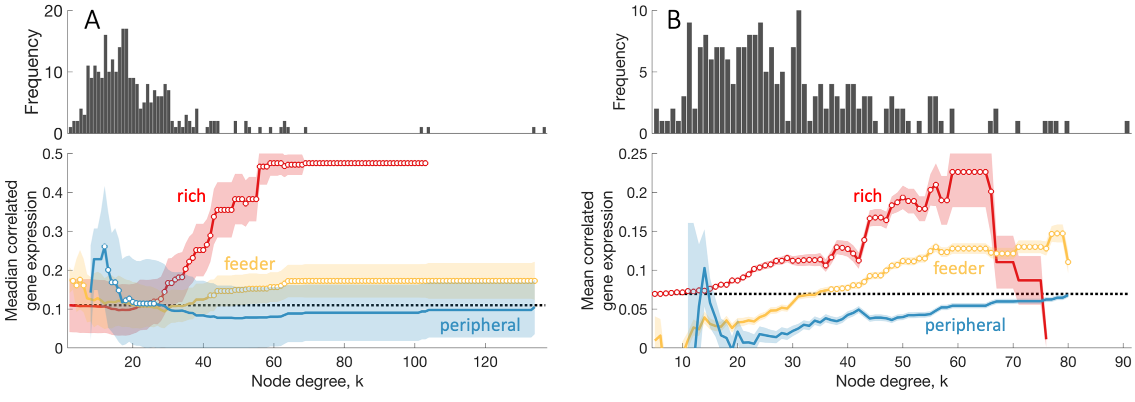


Figure S9: **Consistency of transcriptional coupling results across different species**. **(A)** Median correlated gene expression for rich, feeder and peripheral links as a function of node degree $k$ in the neuronal *C.elegans* connectome. Shaded area corresponds to the half of the median absolute deviation, circles indicate a statistically significant increase in CGE in a given link type relative to the rest of the network (one-sided Wilcoxon rank sum test, $p < 0.05$) [adapted and reproduced from Arnatkevičiūtė et al. [2]]. **(B)** Mean correlated gene expression for rich, feeder, and peripheral links as a function of node degree $k$ in the mouse connectome. Shaded area corresponds to the standard error of the mean, circles indicate a statistically significant increase in CGE for a given link type relative to the rest of the network (one-sided Welch's $t$-test, $p < 0.05$) [adapted and reproduced from Fulcher and Fornito [3]].

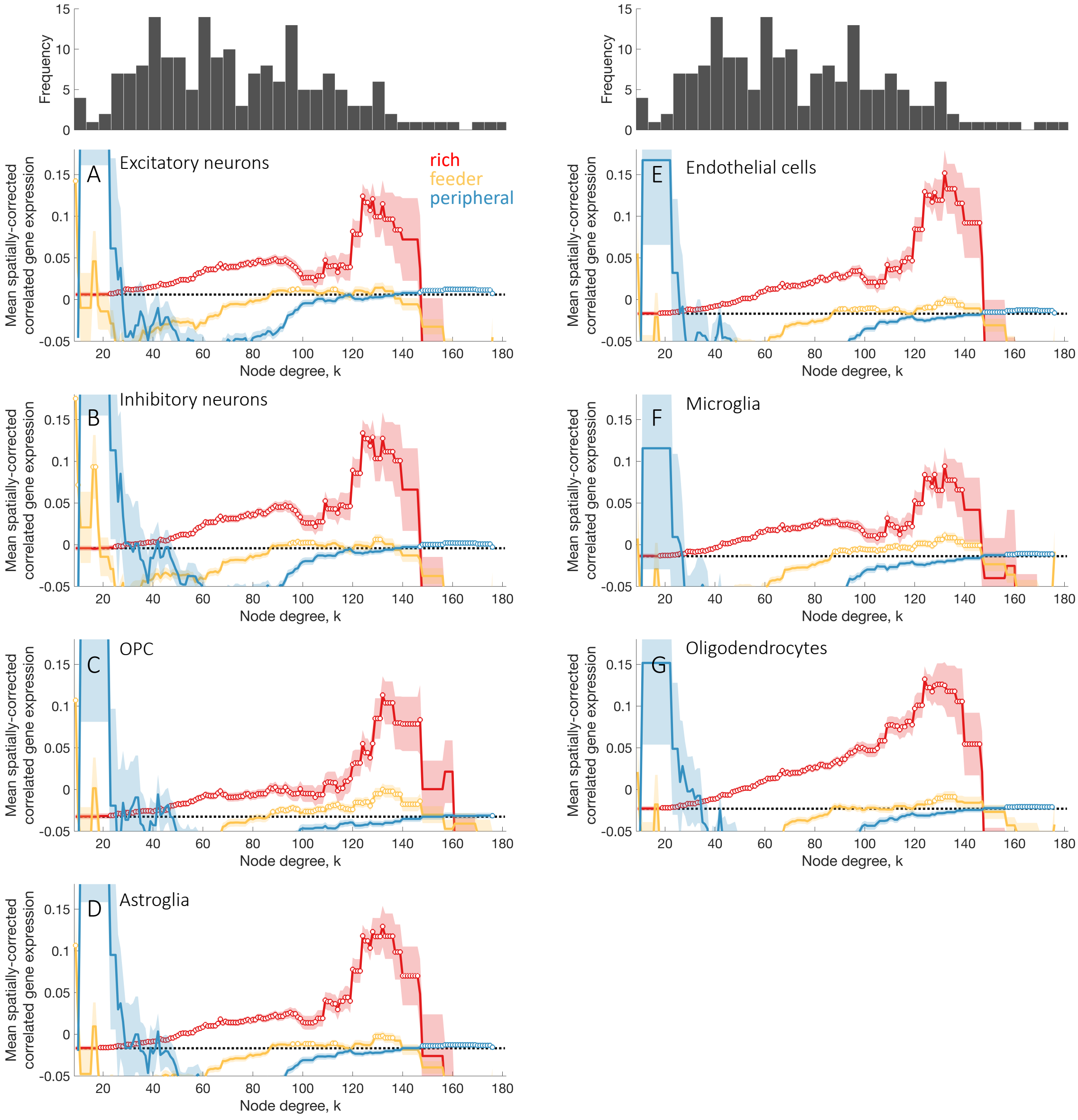


Figure S10: **Cell-specific genes demonstrate elevated transcriptional coupling between connected hub regions.** The top of each panel shows the degree distribution of the representative group-level connectome of brain regions in the left cortical hemisphere. Degree is computed from whole-brain connectivity. The bottom of each panel shows the mean spatially-corrected correlated gene expression of cell-specific genes [excitatory neurons **(A)**, inhibitory neurons **(B)**, oligodendrocyte progenitor cell **(C)**, astroglia **(D)**, endothelial cells **(E)**, microglia **(F)**, oligodendrocytes **(G)**] for rich (hub-hub), feeder (hub-nonhub), peripheral (nonhub-nonhub) connections as a function of degree threshold, $k$ used to define hubs. The mean CGE across all network links shown as a dotted black line. Shaded area corresponds to the standard error of the mean, circles indicate a statistically significant increase in CGE in a given link type compared to the rest of the network (one-sided Welch's $t$-test, $p < 0.05$).

Table S1: **Gene-set enrichment analysis results.** Gene Ontology biological processes categories implicated in the increased transcriptional coupling in rich compared to peripheral links (false discovery rate (FDR) corrected $p < 0.05$).

| Term ID | Description | $p_{\mathrm{FDR}}$ |
|---|---|---|
| GO:0006119 | Oxidative phosphorylation | $1.5 \times 10^{-9}$ |
| GO:0042773 | ATP synthesis coupled electron transport | $1.5 \times 10^{-9}$ |
| GO:0042775 | Mitochondrial ATP synthesis coupled electron transport | $1.5 \times 10^{-9}$ |
| GO:0045333 | Cellular respiration | $1.5 \times 10^{-9}$ |
| GO:0006120 | Mitochondrial electron transport NADH to ubiquinone | $1.4 \times 10^{-3}$ |
| GO:0007007 | Inner mitochondrial membrane organization | $1.4 \times 10^{-3}$ |
| GO:0009145 | Purine nucleoside triphosphate biosynthetic process | $1.4 \times 10^{-3}$ |
| GO:0009206 | Purine ribonucleoside triphosphate biosynthetic process | $1.4 \times 10^{-3}$ |
| GO:0051438 | Regulation of ubiquitin-protein transferase activity | $1.4 \times 10^{-3}$ |
| GO:0009201 | Ribonucleoside triphosphate biosynthetic process | $2.0 \times 10^{-3}$ |
| GO:0022904 | Respiratory electron transport chain | $2.0 \times 10^{-3}$ |
| GO:0042407 | Cristae formation | $2.0 \times 10^{-3}$ |
| GO:0009142 | Nucleoside triphosphate biosynthetic process | $6.2 \times 10^{-3}$ |
| GO:0010257 | NADH dehydrogenase complex assembly | $6.6 \times 10^{-3}$ |
| GO:0032981 | Mitochondrial respiratory chain complex I assembly | $6.6 \times 10^{-3}$ |
| GO:0097031 | Mitochondrial respiratory chain complex I biogenesis | $6.6 \times 10^{-3}$ |
| GO:0051443 | Positive regulation of ubiquitin-protein transferase activity | $1.3 \times 10^{-2}$ |
| GO:0006303 | Double-strand break repair via nonhomologous end joining | $1.3 \times 10^{-2}$ |
| GO:0072655 | Establishment of protein localization to mitochondrion | $1.3 \times 10^{-2}$ |
| GO:0000726 | Non-recombinational repair | $1.6 \times 10^{-2}$ |
| GO:0009060 | Aerobic respiration | $1.7 \times 10^{-2}$ |
| GO:0033108 | Mitochondrial respiratory chain complex assembly | $1.9 \times 10^{-2}$ |
| GO:0006090 | Pyruvate metabolic process | $1.9 \times 10^{-2}$ |
| GO:0070585 | Protein localization to mitochondrion | $1.9 \times 10^{-2}$ |
| GO:0006754 | ATP biosynthetic process | $2.0 \times 10^{-2}$ |
| GO:0000492 | Box C/D snoRNP assembly | $2.0 \times 10^{-2}$ |
| GO:1903146 | regulation of autophagy of mitochondrion | $2.1 \times 10^{-2}$ |
| GO:0006165 | Nucleoside diphosphate phosphorylation | $2.5 \times 10^{-2}$ |
| GO:0046939 | Nucleotide phosphorylation | $2.5 \times 10^{-2}$ |
| GO:1901070 | Guanosine-containing compound biosynthetic process | $2.9 \times 10^{-2}$ |
| GO:0006818 | Hydrogen transport | $3.0 \times 10^{-2}$ |
| GO:0015992 | Proton transport | $3.0 \times 10^{-2}$ |
| GO:1902600 | Hydrogen ion transmembrane transport | $3.0 \times 10^{-2}$ |
| GO:0042776 | Mitochondrial ATP synthesis coupled proton transport | $3.0 \times 10^{-2}$ |
| GO:0006733 | Oxidoreduction coenzyme metabolic process | $3.7 \times 10^{-2}$ |
| GO:0006626 | Protein targeting to mitochondrion | $3.7 \times 10^{-2}$ |
| GO:0061418 | Regulation of transcription from RNA polymerase II promoter in response to hypoxia | $4.6 \times 10^{-2}$ |
| GO:0019362 | Pyridine nucleotide metabolic process | $4.6 \times 10^{-2}$ |
| GO:0046496 | Nicotinamide nucleotide metabolic process | $4.6 \times 10^{-2}$ |
| GO:0006418 | tRNA aminoacylation for protein translation | $4.6 \times 10^{-2}$ |
| GO:0043038 | Amino acid activation | $4.6 \times 10^{-2}$ |
| GO:0043039 | tRNA aminoacylation | $4.6 \times 10^{-2}$ |
| GO:0000470 | maturation of LSU-rRNA | $4.6 \times 10^{-2}$ |
| GO:0006302 | Double-strand break repair | $4.6 \times 10^{-2}$ |
| GO:0006164 | Purine nucleotide biosynthetic process | $4.6 \times 10^{-2}$ |
| GO:0042451 | Purine nucleoside biosynthetic process | $4.6 \times 10^{-2}$ |
| GO:0046129 | Purine ribonucleoside biosynthetic process | $4.6 \times 10^{-2}$ |
| GO:0031146 | SCF-dependent proteasomal ubiquitin-dependent protein catabolic process | $4.7 \times 10^{-2}$ |

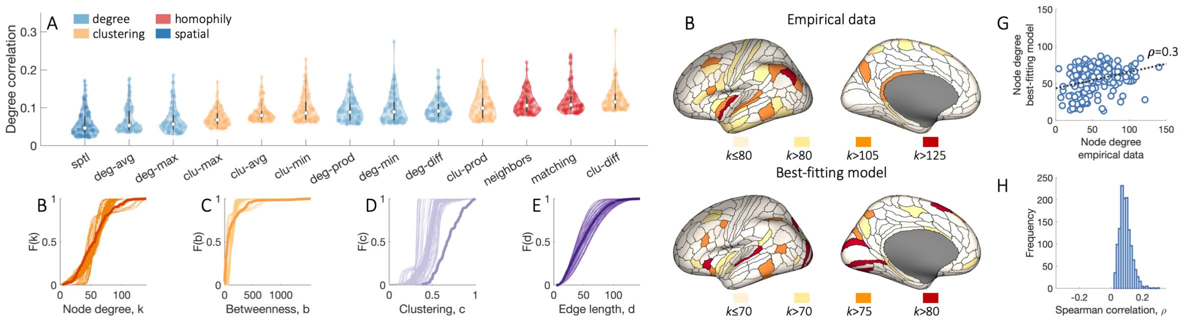


Figure S11: **Generative brain network models optimized to maximise degree sequence correlations using Spearman correlation do not reproduce regional connectivity of the empirical data.** **(A)** Each distribution shows Spearman degree sequence correlations between empirical and model networks within the left hemisphere of the top 100 best-fitting parameter combinations (out of 10 000) for each model, as identified using an optimization procedure explicitly designed to maximize the degree sequence correlation between empirical and model networks. The color of each box indicates the topology metric used in the model: homophily is shown in red, clustering in orange, degree in light blue, and spatial in dark blue. The specific wiring rule names are shown along the horizontal axis. Cumulative distributions of **(B)** node degree, $k$; **(C)** betweenness centrality, $b$; **(D)** clustering coefficient, $c$; and **(E)** edge length, $d$, for the empirical connectome (darker line) and 100 instantiations of the best-fitting 'clu-diff' model corresponding to the data points shown in **A** (lighter lines). Optimization of the degree sequence correlation comes at the expense of capturing topological properties of the connectome, which are not considered in the optimization. **(F)** Spatial distributions of network hubs for the left hemisphere at different levels of $k$ for the empirical data (top) and the best-fitting generative model (bottom). The degree distribution of the model network is very narrow, so different thresholds are applied for visualization. **(G)** Correlation between the degree sequences of the empirical data and the best-fitting model (Spearman $\rho = 0.3$). **(H)** The distribution of correlation values quantifying the relationship of degree sequences within a single hemisphere between empirical data and synthetic networks generated using the top 100 best-fitting parameter combinations for each of the 13 generative models, corresponding to the data points in **A**.
